# Quantitative imaging of native tissue architecture combined with deep multi-omics reveals the spatial biology of kidney aging

**DOI:** 10.64898/2026.10.28.762545

**Authors:** William V. Trim, Mariia Diakova, Matthew Sonnett, Kseniia Petrova, Bingsen Zhang, Seungeun Oh, Romain Donne, Corissa McEwen, Hunter Salmon, Abhilash Gadi, Tea Shavlakadze, Edward T. Chouchani, David J. Glass, Marc W. Kirschner

## Abstract

Aging alters tissue architecture and metabolism in spatially heterogeneous ways, yet the connections between these measurable structural and compositional phenotypes and underlying molecular changes are largely unknown. Here, we introduce NORIGAMI (NoRI-Guided AI for Multimodal Integration), a pipeline combining Normalized Raman Imaging (NoRI), deep learning, quantitative morphology, deep proteomics, and metabolomics to investigate renal aging. Unlike conventional histology, NoRI provides label-free, scattering-corrected measurements of absolute total protein and lipid mass concentrations (mg/mL). In 32 male rats aged 9, 18, 21 and 25 months, NORIGAMI predicted the chronological age of held-out rats from NoRI images (mean absolute error 0.78 months; R² = 0.97), and whole-tissue images outperformed models trained on isolated tubules. Quantitative imaging identified 41 age-associated morphological and compositional features (FDR < 0.05), revealing anatomically heterogeneous and non-linear changes in lipid droplet concentration, droplet size, cytoplasmic lipid concentration and tubular architecture. Deep molecular profiling quantified 11,485 proteins and 269 metabolites, revealing coordinated declines in differentiated proximal tubule (PT) identity and oxidative metabolism, selective increases in basement-membrane collagen-IV chains, and increasing molecular heterogeneity with age. A protein clock, built mainly from extracellular-matrix and heparan-sulfate proteins, predicted age with a mean absolute error of 1.6 months. Old kidneys diverged along a dominant injury-remodeling axis, seen separately in the proteome, in kidney RNA from 54 other rats and in the metabolome, in which tubular-injury and immune proteins, including the NAD^+^-consuming enzyme CD38, rise as differentiated-tubule proteins and NAD^+^ fall. NORIGAMI subsequently linked NAD^+^-associated molecular changes to candidate quantitative features of tubular geometry and protein–lipid organization in exploratory analyses, while complementary KIM-1-centered analyses identified overlapping candidate morphological associations involving epithelial injury and, possibly, lipid handling and yielded additional molecular candidates for mechanistic validation. Collectively, these findings suggest that declining tubular metabolic function, altered lipid organization and injury-associated molecular programs represent interconnected features of renal aging. NORIGAMI provides a framework for identifying the quantitative structural manifestations of molecular aging and generating spatially informed, experimentally testable hypotheses about tissue dysfunction.

## Introduction

Aging represents a progressive decline in cellular, tissue, and whole-body physiological function, resulting in a progressive loss of homeostasis, leading to increased vulnerability to chronic disease (Niccoli & Partridge, 2012). Characteristics of aging include genomic instability, loss of proteostasis, altered nutrient sensing, mitochondrial dysfunction, cellular senescence, and chronic low-grade inflammation (López-Otín et al., 2013; López-Otín et al., 2023). These features are frequently measured in isolation in cell culture or from bulk measurements in tissues, but they collectively converge on energy metabolism and inflammation. Consistent with this, the most profound life-extending interventions (calorie restriction, genetic/pharmacological modulation) are ones that directly influence such systems (Fontana et al., 2010). However, the mechanisms of aging within tissues, and their spatial variability, are poorly understood. Not all cells within a tissue exhibit the same basal metabolism, and inflammatory signals are often localized to regions of specific damage. Aging is therefore unlikely to be uniform across a tissue. Indeed, an understanding of the circuitry and spatiality of tissue homeostasis is increasingly important for understanding tissue health and disease (Meizlish et al., 2021). Conventional biochemical methods such as transcriptomics, proteomics, and metabolomics are rich in information but do not address changes to the native organization of tissues. Linking molecular changes to quantitative changes in tissue composition with age remains an unmet need.

The kidney in many ways is a compelling organ to address this gap. It is a highly energetic organ with striking metabolic heterogeneity. The proximal tubules rely heavily on fatty acid oxidation to power solute reabsorption, while the distal tubules and collecting ducts are more glycolytic and serve to maintain homeostasis (Bhargava & Schnellmann, 2017; Kang et al., 2015). Due to its limited regenerative capacity, the kidney is acutely sensitive to mitochondrial and proteostatic perturbations, with lipid accumulation representing a primary pathological feature of kidney injury (Giardini et al., 2024; Jang et al., 2020; Młynarska et al., 2026; Ren et al., 2023). The prevalence of chronic kidney disease rises sharply with age. Structurally, renal aging manifests as nephron loss, glomerular enlargement, glomerular basement membrane (GBM) thickening, mesangial expansion and glomerulosclerosis, tubular atrophy, interstitial fibrosis and microvascular rarefaction. These changes are strikingly conserved across species (Denic et al., 2016); in aging rats they overlap with chronic progressive nephropathy, a common rat-specific disease (Hard & Khan, 2004). In the rat, the trajectory of age-related kidney disease has recently been charted at the transcript level across 6–27 months of age (Shavlakadze et al., 2026), revealing sustained induction of immune–inflammatory and extracellular-matrix remodeling programs.

The role of lipids in renal aging and disease appears to be important. When fatty acid oxidation is impaired, excess lipid accumulates in droplets, driving endoplasmic reticulum stress, oxidative damage, inflammation, and fibrosis (Kang et al., 2015). However, these processes have not been studied quantitatively in aging kidneys. Tissue nicotinamide adenine dinucleotide (NAD^+^) levels decline with age (Camacho-Pereira et al., 2016; Covarrubias et al., 2021) and, given the role of NAD^+^ in sustaining fatty acid oxidation, this decline may provide at least a partial explanation for lipid accumulation. NAD^+^ also acts as a substrate for sirtuins and PARPs, which govern mitochondrial energetics, stress responses and DNA repair/damage (Covarrubias et al., 2020; Covarrubias et al., 2021). Consistent with these findings, kidney injury in humans is accompanied by impaired *de novo* NAD^+^ biosynthesis (Poyan Mehr et al., 2018), while supplementation with an NAD^+^ precursor protects aged mice against acute kidney injury (Guan et al., 2017). Levels of CD38, a major NAD^+^-consuming NADase that cleaves NAD^+^ into nicotinamide and ADP-ribose, increase substantially with age, and CD38 deletion prevents age-related NAD^+^ decline (Camacho-Pereira et al., 2016). These observations led us to explore whether changes in kidney morphology are linked to increased NAD^+^ consumption by CD38.

Connecting the molecular circuitry of aging in the kidney to structural topography features requires simultaneous quantitative spatially resolved and molecular measurements. Conventional histology, the cornerstone of renal pathology, relies on fixation, embedding and staining, procedures that distort tissue architecture, extract lipid and introduce batch-to-batch variability, limiting quantitative comparisons and high-resolution machine-learning analyses. Fluorescence imaging of fixed tissues preserves spatial context but requires specific labels and therefore reports only on pre-selected targets. Bulk omics provide molecular depth but lose the spatial heterogeneity that characterizes tissues as structurally complex as the kidney.

We therefore turned to Normalized Raman Imaging (NoRI), developed in our laboratory, which overcomes some of the conventional imaging constraints by normalizing protein and lipid signals against the combined protein, lipid and water signal. This largely removes the effects of scattering and yields absolute local concentrations (mg/mL) of proteins and lipids at a resolution of 0.4 µm voxels. Measurements remain quantitative up to 0.5 mm into tissue and are directly comparable across specimens (Oh et al., 2022). Machine-learning methods can extract subtle but useful information even from noisy sources and should benefit further from calibrated, quantitative inputs. We recently used NoRI to explore reperfusion injury in the kidney (Trim, Oh et al., 2026, in press), showing that the quantitative output of NoRI supports automated analysis by convolutional neural networks (CNNs). Tubule segment, anatomical region and sex could be identified directly from NoRI data with high accuracy, and NoRI allowed us to characterize quantitatively the reorganization of protein and lipid during recovery from hypoxia. Applied to aging, this approach offers a data-driven complement to curated hallmark checklists. CNNs can learn the features, known and unknown, that distinguish kidneys of different ages, while NoRI assigns each an absolute magnitude of change across the lifespan.

In this report, we set out to study the physiology of kidney aging in the rat using NoRI microscopy, proteomics and metabolomics, measured in the paired kidneys of each rat: one kidney was imaged, and the contralateral kidney was used for proteomics and metabolomics. To do this we introduced a CNN-based pipeline called NoRI-Guided AI for Multimodal Integration (NORIGAMI). NORIGAMI co-aligns age-associated NoRI features with protein and metabolite abundances, relating molecular changes to the specific tissue structural features and compositional lesions with which they co-vary. Using the NORIGAMI pipeline, we demonstrate a strong predictive ability for chronological age and quantify the absolute magnitude and spatial distribution for each feature identified as correlating with aging. By co-aligning these features with proteomic and metabolomic profiles from the same animals, we look for the molecular programs that accompany these morphological features. Among these, proteins and metabolites of NAD^+^ synthesis and consumption, including CD38, were associated with specific morphological features of aging in exploratory analyses, pointing to altered NAD^+^ handling as a candidate link between molecular and structural aging. Together, this work moves beyond a qualitative hallmark framework toward a quantitative, spatially grounded and molecularly resolved account of how the kidney’s metabolic architecture is remodeled from adulthood into old age in the male rat. More generally, NORIGAMI can relate quantitative morphological features to proteomic, metabolomic or other numerical datasets measured in the same animals.

## Results

Our overall data collection and analysis approach is shown in **Figure 1**. We measured native tissue architecture by NoRI and related it to protein and metabolite levels in the normally aging kidney, using four ages with eight rats per age. We first combined NoRI with feature segmentation and automated histopathology to create a quantitative morphological atlas of the kidney across ages (**Figure 1A**). The features we identify allow the age of a kidney specimen to be predicted from NoRI data alone. We then collected bulk-tissue proteomics and metabolomics data from the contralateral kidneys of the same animals. Next, we integrated these data types using NORIGAMI, a multi-layer neural network that uses variations in individual protein or metabolite levels in different animals of the same age to connect changes in NoRl-identified spatial features to protein pathways and metabolites (**Figure 1B,C**; **Supplementary Figure 1**; Methods). Candidate proteins can then be stained with antibodies on the imaged specimens and overlaid on the NoRI images (**Figure 1A** shows KIM-1 as an example). By this means, NORIGAMI provides a new framework for visualizing kidney aging.

**Figure 1.**
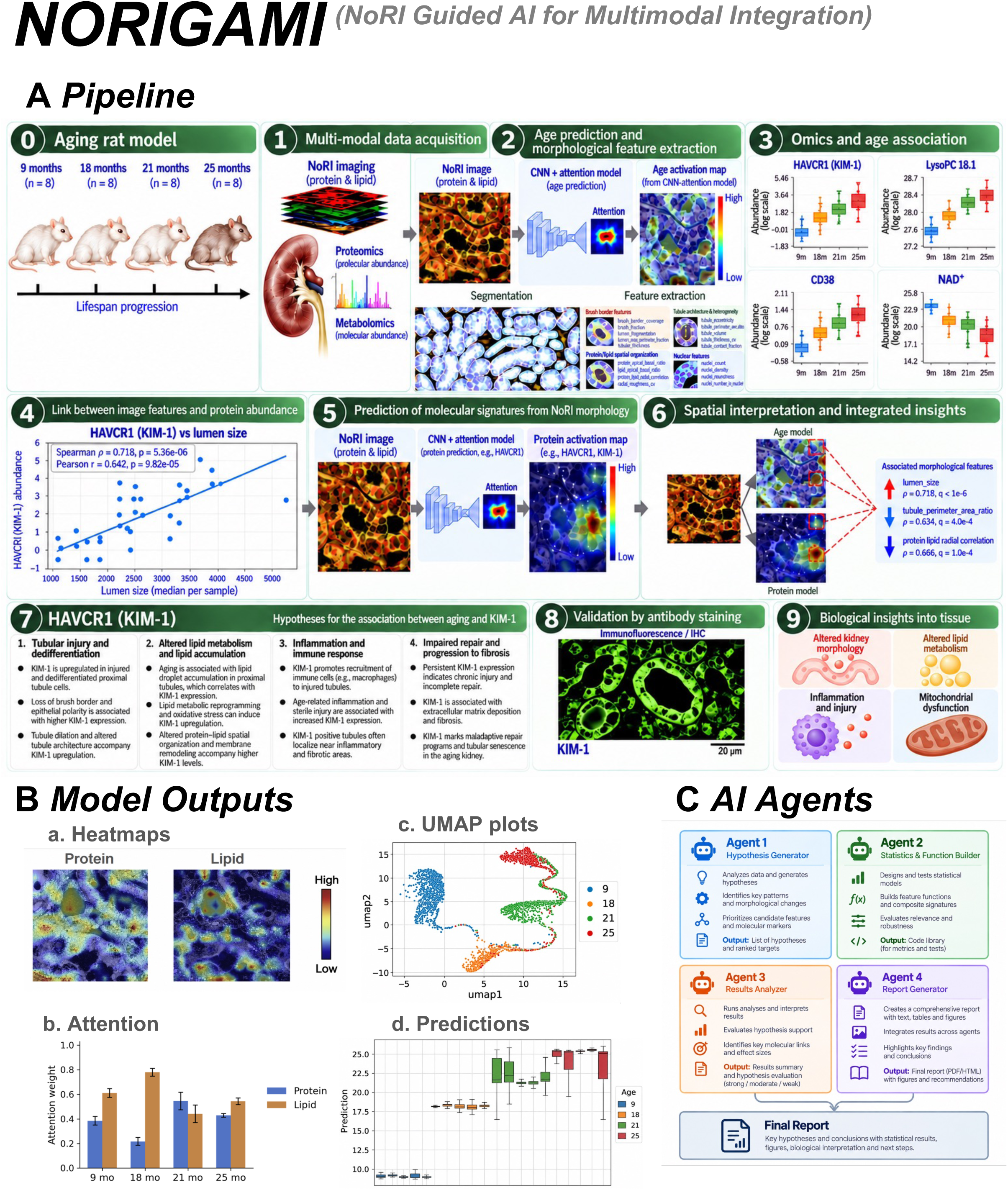
NORIGAMI links NoRI imaging, tissue morphology, proteomics and metabolomics to study kidney aging. (A) The NORlGAMl pipeline. Kidneys were collected from 32 rats (n = 8 per age) at 9, 18, 21 and 25 months; one kidney from each rat was imaged by Normalized Raman lmaging (NoRl) and the other was used for proteomics and metabolomics. The pipeline predicts age from NoRl images, extracts morphological features, finds associations between molecules and morphology, predicts molecular abundance from tissue images and maps these predictions back onto the tissue. Selected candidates, such as KlM-1 (HAVCR1), are followed up by antibody staining. (B) Example model outputs: (a) protein and lipid attention maps showing the image regions that drive predictions; (b) relative contribution of the protein and lipid channels at each age; (c) learned image representations, arranged by age; (d) model age estimates for each age group. (C) Al-assisted interpretation. A multi-agent framework combines morphological features, age predictions and molecular measurements to propose testable hypotheses about tissue aging.

We first tested whether NoRI images alone could predict age. We trained ten CNN models, each with training and validation animals kept separate and each validated on four held-out rats, one from each age group (Methods). The models scored individual NoRl image tiles (320 × 320 pixels; 132.5 × 132.5 µm), and tile predictions were summarized for each rat as their median. A UMAP of the learned tile representations largely separated the four ages (**Figure 2A**), and **Figure 2B** shows each tile’s predicted age on the same map. Across the ten models, the median animal-level mean absolute error (MAE) was 0.77 months, compared with 7.06 months when age labels were shuffled among the validation rats (**Supplementary Figure 2**). Pooling the predictions for each held-out rat across models gave an MAE of 0.78 months (95% bootstrap interval 0.57–1.00) and R² = 0.97 (95% interval 0.96–0.99; **Figure 2C**). Tile predictions varied within and between kidneys. Within-animal variability, the mean standard deviation of tile predictions, was lowest at 18 months (1.03 months), compared with 2.20 months at 9 months, 2.03 months at 21 months and 2.44 months at 25 months. Kidneys differed more from one another at 21 and 25 months than at 9 and 18 months (**Figure 2C**), and predictions also varied within individual kidneys.

**Figure 2.**
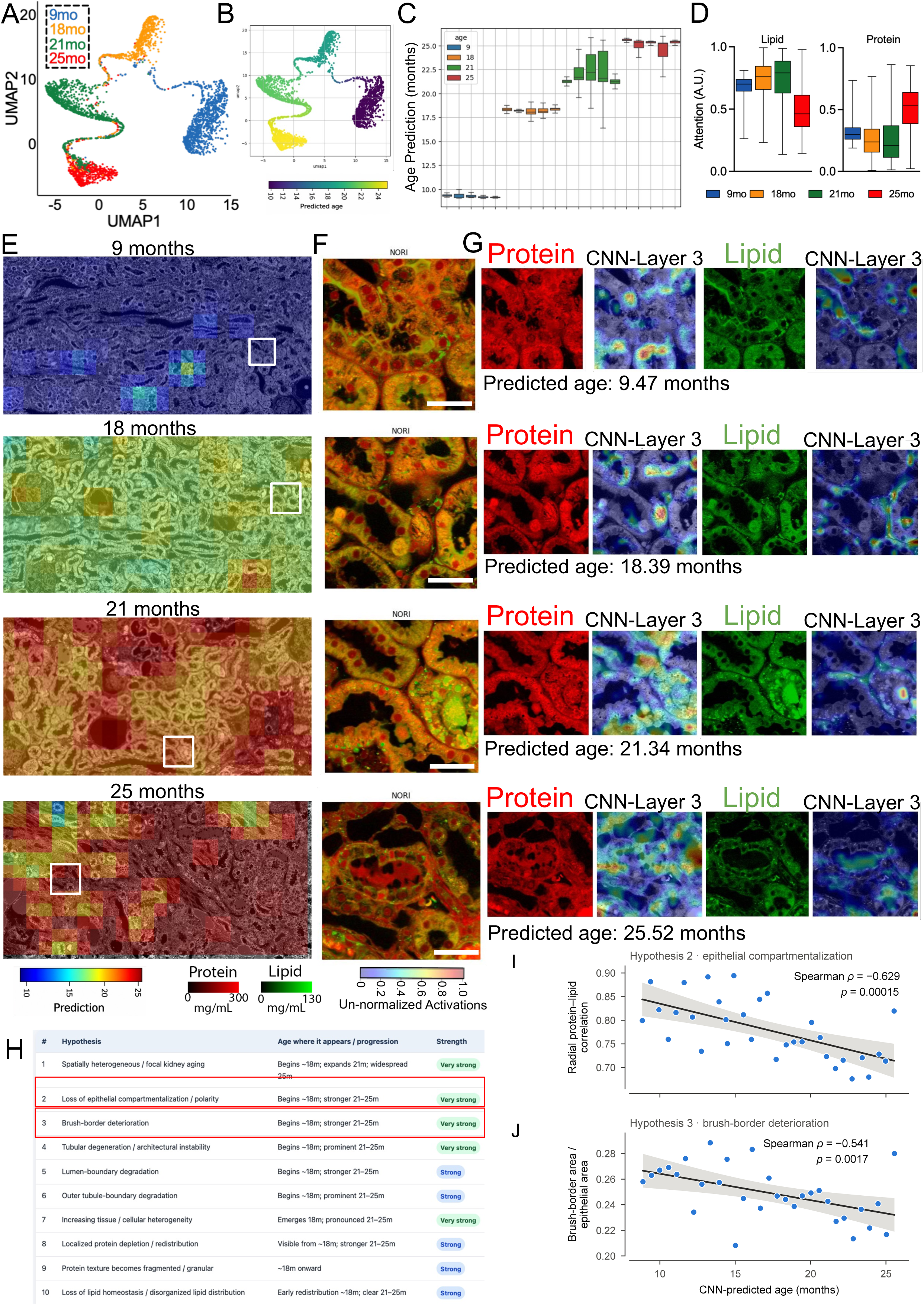
NoRI images predict age, and agentic AI proposes candidate age-associated features at subcellular resolution. (A) UMAP of the age model’s learned representations of NoRI image tiles (320 × 320 pixels; 132.5 × 132.5 µm) from held-out rats (n = 5 per group) aged 9, 18, 21 and 25 months, colored by age group. (B) The same UMAP colored by predicted age. (C) Predicted age for held-out rats that were not used in training. Each box summarizes all tiles from one animal (n = 5 held-out rats per age). (D) Relative attention given by the model to the quantitative lipid and protein channels at each age. (E) Whole-section NoRI images from rats aged 9, 18, 21 and 25 months (top to bottom), with each tile colored by its predicted age. White boxes mark the tiles shown in F and G. (F) NoRI composite of each boxed tile (protein, red; lipid, green). (G) Protein and lipid concentration maps, each with its CNN layer-3 activation map, which highlights the regions that contribute most to the prediction; the tile’s predicted age is shown below. (H) Ranked hypotheses for the image features that best separate age groups, generated by the NORIGAMI agentic framework (GPT-4-based agents) from the age model’s outputs and segmented NoRI structures. The table gives the age at which each feature appears, how it progresses and the strength of support. (I, J) Two of these features measured directly and plotted against predicted age: radial protein–lipid correlation across the tubular epithelium (I; hypothesis 2) and brush-border area relative to epithelial area (J; hypothesis 3). In C and D, boxes show medians and interquartile ranges of tile values. In I and J, tiles from all rats were grouped into 0.5-month bins of predicted age. Each point is one bin, lines are linear fits with 95% confidence bands, and Spearman’s p is shown with its two-sided *P* value (t approximation with n – 2 degrees of freedom, where n is the number of bins). Scale bars, 40 µm.

We next asked how much the protein and lipid channels each contribute to age prediction. **Figure 2D** shows the relative attention given to the lipid and protein channels across five validation rats per age group, revealing the considerable contribution of quantitative lipid information for predicting age in 9-, 18-, and 21-month rats (mean attention 0.68–0.74 for lipid and 0.26–0.32 for protein). lmportantly, standard histological processing extracts much of the tissue lipid, so this observation would not be available from conventional paraffin-embedded stained sections. By 25 months, attention was more evenly distributed between the two modalities (0.50 ± 0.19 for both protein and lipid; **Figure 2D**).

We then mapped predicted age across tissue sections. **Figure 2E,F** provide examples of features of kidney aging. For one representative kidney section per age group, we overlaid tile-level predicted age (132.5 × 132.5 µm tiles; color scale at the bottom of panel E) onto the original NoRI image. From each large-area image we extracted one tile (white box in **Figure 2E**, displayed in **Figure 2F**). Each extracted NoRI tile was chosen as an example of its age group. **Figure 2G** shows the quantitative protein (red) and lipid (green) channels of each tile, the corresponding CNN layer 3 activation maps and the tile’s predicted age. Together, these maps suggest candidate features of kidney aging at both tissue and subcellular scales in minimally processed, fixed tissue.

NORIGAMI also includes an AI-agent framework that proposes candidate image features from the model outputs (**Figure 1C**). **Figure 2H** lists the ten candidate age-associated features it proposed. Combined with automated segmentation of tubules, nuclei, brush border and other structures, such features can be measured in each NoRI image tile and compared with predicted age. **Figure 2I,J** show two of these proposed features measured directly by automated segmentation of NoRl images. Both correlated with CNN-predicted age across bins of image tiles, linking interpretable morphological characteristics to spatial patterns of predicted kidney aging.

To investigate the contribution of tissue context to age prediction, we compared CNN models for NoRl tiles (320 × 320 pixels) with individually segmented tubules (256 × 256 pixels). Although isolated proximal tubules contained substantial age-associated information, their predictions separated successive age groups less well than those from full-tissue tiles (**Figure 3A**). **Figure 3B,C** compare the attention given to the protein and lipid channels by the whole-tile and tubule models. Aggregating predictions from multiple individual tubules reduced variability but did not eliminate a systematic overestimation of age in young animals (not shown). These data suggest that full-tissue tiles provide a stronger representation of kidney aging, possibly through integrating morphological and chemical information across multiple tubules and their surrounding tissue. Consistent with this interpretation, quantitative analysis identified age-associated changes in proximal tubule connection regions, which CNN attribution maps also appeared to highlight (not shown). Together, these observations suggest that tissue-level spatial organization, instead of isolated tubular morphology alone, may contribute to accurate kidney age prediction.

**Figure 3.**
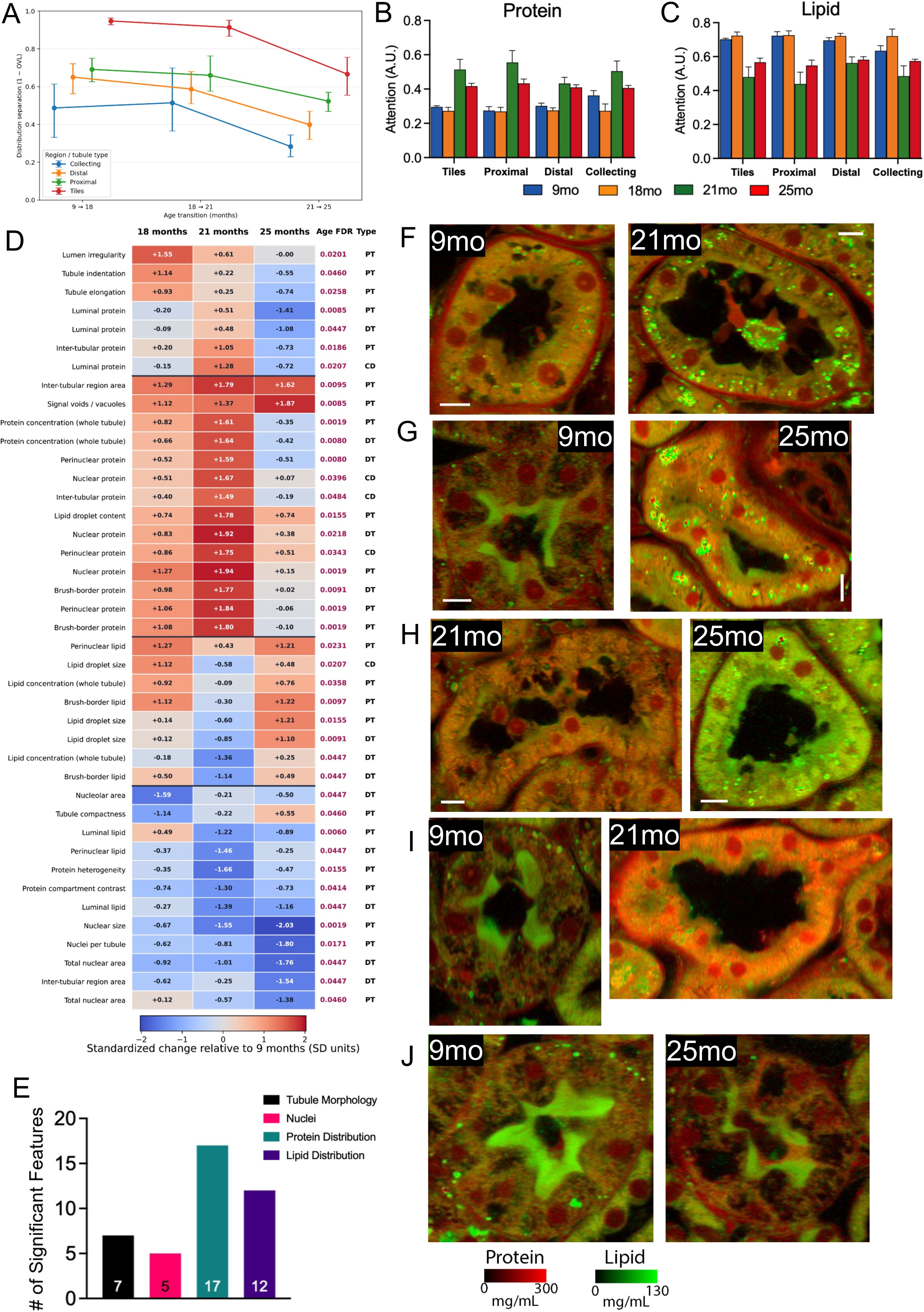
Age-associated changes in tubular protein, lipid and morphology. (A) Separation between successive age groups (1 – overlap coefficient) for age-prediction models trained on whole tiles or on segmented tubules. (B, C) Relative attention given to the protein (B) and lipid (C) channels at each age by the whole-tile and tubule models. (D) The 41 features that change with age, measured in proximal tubules (PT), distal tubules (DT) and collecting ducts (CD). Values are changes from 9 months in standard-deviation units (red, increase; blue, decrease). Each feature was summarized per animal (median) and tested across the four ages with the Kruskal–Wallis test, with Benjamini–Hochberg correction within each tubule type; all features shown have FDR < 0.05 (adjusted *P* values at right). (E) The 41 features grouped by type: tubular morphology (7), nuclear organization (5), protein distribution (17) and lipid measurements (12). (F–J) Example NoRI images of tubules with low and high values (left and right) for lipid droplet concentration (F), lipid droplet size (G), cytoplasmic lipid concentration (H) and cytoplasmic protein concentration (I), and with high and low brush-border lipid concentration (J). Absolute concentration scales are presented below J. A, mean ± 95% CI; B and C, mean ± SEM. Analyses in D and E use per-animal values (up to n = 8 per age). Significance for pairwise comparisons was defined as BH-adjusted P < 0.05. Scale bars, 10 µm.

Using NORlGAMl, we measured 51 morphological, protein and lipid features of each segmented tubule and summarized them separately for proximal tubules, distal tubules and collecting ducts, giving 153 features excluding glomerular features which are not presented in the present analysis (feature definitions in **Supplementary Table 1**). Each feature was summarized per animal and tested across the four ages with the Kruskal–Wallis test, with Benjamini–Hochberg correction within each tubule type. Forty-one features changed with age (FDR < 0.05; **Figure 3D**): 17 describing protein distribution, 12 lipid measurements, 7 tubular morphology and architecture, and 5 nuclear organization (**Figure 3E**). Because NoRl measures total lipid concentration in minimally processed tissue, these lipid concentrations cannot be obtained in absolute units from conventional histology.

**Figure 3F–J** shows example tubules for five features that NoRl measures in fixed tissue without lipid extraction or staining: lipid droplet concentration and size, cytoplasmic lipid and protein concentration, and brush-border lipid concentration. Several of the age changes were not monotonic. Lipid droplet concentration in proximal tubules rose most at 21 months, protein concentrations peaked at 21 months, and lipid droplet size fell at 21 months before rising at 25 months (**Figure 3D**). Because each age group was collected on a single day, a batch contribution to these non-monotonic changes, particularly the 21-month rise in protein concentration across compartments, cannot be excluded. **Table 1** lists values and statistics for each age group for 25 of the protein and lipid features in **Figure 3D**, including four of the five shown in **Figure 3F–J**.

**Table 1.** Age-associated changes in quantitative morphological features of kidney tubules measured by NoRI across aging (*n* =8/ group). Statistical differences between age groups were assessed using Kruskal–Wallis.

| **Feature identifiers** |  | **Molecular Content (mg/mL) mean ± SD (rat means)** |  |  |  | **Kruskal–Wallis (rat medians)** |  |  |  | **Mann–Whitney U post hoc: six BH-adjusted P values** |  |  |  |  |  |
|---|---|---|---|---|---|---|---|---|---|---|---|---|---|---|---|
| **Feature name** | **Tubule type** | **9 months** | **18 months** | **21 months** | **25 months** | **H (χ<sup>2</sup>)** | **df** | **KW P** | **KW BH FDR** | **9 vs 18** | **9 vs 21** | **9 vs 25** | **18 vs 21** | **18 vs 25** | **21 vs 25** |
| **Luminal protein** | PT | 38.72 ± 5.05 | 38.44 ± 7.13 | 43.42 ± 4.56 | 29.24 ± 3.96 | 15.72 | 3 | **0.001** | 0.009 | 0.798 | 0.281 | **0.002** | 0.241 | **0.030** | **<0.001** |
| **Luminal protein** | DT | 36.08 ± 8.86 | 36.36 ± 9.56 | 41.83 ± 3.04 | 28.81 ± 6.49 | 11.17 | 3 | **0.011** | 0.045 | 0.878 | 0.688 | 0.076 | 0.157 | 0.076 | **0.004** |
| **Inter-tubular protein** | PT | 69.24 ± 14.63 | 71.67 ± 10.32 | 82.99 ± 10.75 | 58.99 ± 6.17 | 12.51 | 3 | **0.006** | 0.019 | 0.645 | 0.084 | 0.234 | 0.157 | 0.157 | **<0.001** |
| **Luminal protein** | CD | 17.45 ± 2.09 | 17.18 ± 2.22 | 23.54 ± 3.76 | 15.18 ± 1.93 | 16.87 | 3 | **<0.001** | 0.021 | 0.721 | **0.021** | **0.042** | **0.009** | 0.126 | **<0.001** |
| **Protein concentration (whole tubule)** | PT | 110.51 ± 9.66 | 128.33 ± 9.19 | 142.46 ± 13.75 | 106.96 ± 14.90 | 19.68 | 3 | **<0.001** | 0.002 | **0.010** | **<0.001** | 0.328 | 0.060 | **0.009** | **0.003** |
| **Protein concentration (whole tubule)** | DT | 99.37 ± 7.73 | 108.04 ± 9.41 | 119.14 ± 7.00 | 96.23 ± 6.41 | 18.58 | 3 | **<0.001** | 0.008 | 0.100 | **<0.001** | 0.234 | 0.100 | **0.014** | **<0.001** |
| **Perinuclear protein** | DT | 106.19 ± 6.51 | 114.69 ± 10.33 | 127.34 ± 7.28 | 101.90 ± 8.07 | 19.15 | 3 | **<0.001** | 0.008 | 0.161 | **<0.001** | 0.156 | **0.031** | **0.030** | **<0.001** |
| **Nuclear protein** | CD | 77.97 ± 3.68 | 84.89 ± 11.94 | 99.97 ± 16.89 | 79.81 ± 3.54 | 13.73 | 3 | **0.003** | 0.040 | 0.688 | **0.003** | 0.798 | **0.030** | 0.688 | **0.002** |
| **Inter-tubular protein** | CD | 49.01 ± 8.22 | 54.02 ± 14.53 | 68.29 ± 8.18 | 47.29 ± 8.15 | 12.82 | 3 | **0.005** | 0.048 | 0.721 | **0.003** | 0.721 | 0.100 | 0.663 | **0.002** |
| **Lipid droplet content** | PT | 69.46 ± 11.14 | 79.35 ± 9.58 | 93.88 ± 11.64 | 81.46 ± 12.88 | 13.21 | 3 | **0.004** | 0.015 | 0.078 | **0.007** | 0.078 | 0.062 | 0.878 | 0.076 |
| **Nuclear protein** | DT | 80.51 ± 7.25 | 91.66 ± 10.88 | 104.52 ± 5.99 | 86.65 ± 9.04 | 14.53 | 3 | **0.002** | 0.022 | 0.196 | **<0.001** | 0.721 | **0.030** | 0.606 | **0.009** |
| **Perinuclear protein** | CD | 73.82 ± 4.23 | 82.66 ± 12.11 | 94.73 ± 13.42 | 79.51 ± 6.38 | 14.64 | 3 | **0.002** | 0.034 | 0.075 | **<0.001** | 0.075 | 0.078 | 0.721 | 0.062 |
| **Nuclear protein** | PT | 98.00 ± 7.81 | 124.93 ± 7.26 | 138.84 ± 17.73 | 102.14 ± 13.88 | 21.32 | 3 | **<0.001** | 0.002 | **<0.001** | **<0.001** | 0.645 | 0.193 | **0.004** | **0.002** |
| **Brush-border protein** | DT | 90.17 ± 5.26 | 101.17 ± 9.89 | 108.91 ± 7.02 | 91.79 ± 8.24 | 16.85 | 3 | **<0.001** | 0.009 | **0.030** | **0.002** | 0.878 | 0.078 | 0.075 | **0.003** |
| **Perinuclear protein** | PT | 112.49 ± 9.41 | 133.64 ± 9.03 | 148.87 ± 14.41 | 113.80 ± 14.24 | 19.99 | 3 | **<0.001** | 0.002 | **0.006** | **<0.001** | 1.000 | 0.126 | **0.007** | **0.002** |
| **Brush-border protein** | PT | 102.86 ± 6.31 | 119.04 ± 6.64 | 130.30 ± 12.25 | 101.41 ± 9.74 | 21.00 | 3 | **<0.001** | 0.002 | **<0.001** | **<0.001** | 0.574 | 0.156 | **0.007** | **0.001** |
| **Perinuclear lipid** | PT | 36.94 ± 3.98 | 44.70 ± 3.76 | 39.04 ± 3.45 | 44.43 ± 8.02 | 11.78 | 3 | **0.008** | 0.023 | **0.007** | 0.281 | 0.076 | **0.044** | 0.959 | 0.196 |
| **Lipid concentration (whole tubule)** | PT | 36.48 ± 4.22 | 42.53 ± 3.22 | 36.79 ± 2.83 | 41.81 ± 7.79 | 10.59 | 3 | **0.014** | 0.036 | **0.009** | 0.959 | 0.241 | **0.009** | 0.959 | 0.241 |
| **Brush-border lipid** | PT | 49.55 ± 2.76 | 53.69 ± 2.33 | 46.49 ± 3.21 | 53.36 ± 4.83 | 14.77 | 3 | **0.002** | 0.010 | **0.009** | 0.281 | **0.042** | **0.009** | 0.798 | **0.021** |
| **Lipid concentration (whole tubule)** | DT | 43.01 ± 3.58 | 42.81 ± 3.79 | 37.08 ± 2.76 | 44.61 ± 4.72 | 11.79 | 3 | **0.008** | 0.045 | 0.721 | **0.014** | 0.721 | **0.014** | 0.492 | **0.014** |
| **Brush-border lipid** | DT | 46.36 ± 3.77 | 49.38 ± 5.19 | 41.08 ± 4.25 | 49.92 ± 3.80 | 11.71 | 3 | **0.008** | 0.045 | 0.459 | 0.076 | 0.418 | **0.009** | 0.798 | **0.009** |
| **Luminal lipid** | PT | 13.24 ± 1.38 | 14.42 ± 1.54 | 11.30 ± 1.52 | 11.63 ± 1.75 | 16.87 | 3 | **<0.001** | 0.006 | 0.382 | **0.003** | 0.075 | **0.003** | **0.009** | 0.281 |
| **Perinuclear lipid** | DT | 48.09 ± 4.13 | 46.79 ± 4.32 | 40.90 ± 3.30 | 47.27 ± 6.25 | 10.93 | 3 | **0.012** | 0.045 | 0.758 | **0.007** | 1.000 | **0.021** | 1.000 | 0.056 |
| **Protein heterogeneity** | PT | 0.15 ± 0.01 | 0.15 ± 0.01 | 0.13 ± 0.01 | 0.14 ± 0.02 | 13.52 | 3 | **0.004** | 0.015 | 0.394 | **0.007** | 0.241 | **0.009** | 0.721 | **0.030** |
| **Luminal lipid** | DT | 13.19 ± 2.81 | 12.51 ± 1.96 | 9.41 ± 0.76 | 10.95 ± 2.77 | 11.93 | 3 | **0.008** | 0.045 | 0.798 | **0.007** | 0.100 | **0.014** | 0.124 | 0.798 |

*Abbreviations: BH, Benjamini–Hochberg; CD, collecting duct; DT, distal tubule; FDR, false-discovery rate; KW, Kruskal-Wallis; PT, proximal tubule; SD, standard deviation.*

### A deep proteomic and metabolomic atlas of the aging rat kidney

To probe the molecular circuitry of aging in the rat kidney, we performed proteomics and metabolomics on the kidney contralateral to the one used for NoRl imaging. Proteomics was performed on 31 rats (seven at 9 months and eight at each of 18, 21 and 25 months) and metabolomics on 31 rats (eight, eight, seven and eight), 30 of which were measured by both. A total of 269 annotated metabolites were quantified across all kidneys (**Supplementary Table 3**). Deep and quantitative proteomic coverage was achieved using a TMT-MS3 method by analysis of 23 high-pH reversed-phase fractions where each sample was analyzed by coupling long 6.5-hour HPLC gradients, 4 FAlMS compensation voltages (Schweppe et al., 2019), and MS2 scans that were performed in the Orbitrap as opposed to most MS3 methods that use the ion trap. Of 11,744 proteins identified at a 1% protein-level FDR, 11,485 were quantified in all 31 animals (**Supplementary Table 2**). Our proteomic depth exceeds that of three recent high-quality whole-kidney aging proteomes in mouse (Keele et al., 2025; Wang et al., 2025; Takasugi et al., 2024), which quantified 9,058, 5,697, and 5,165 proteins, respectively, in every animal (**Figure 4A**). The added depth greatly increased coverage of low abundance protein classes such as transcription factors that are often sparsely represented in tissue proteomes (**Figure 4A**).

**Figure 4.**
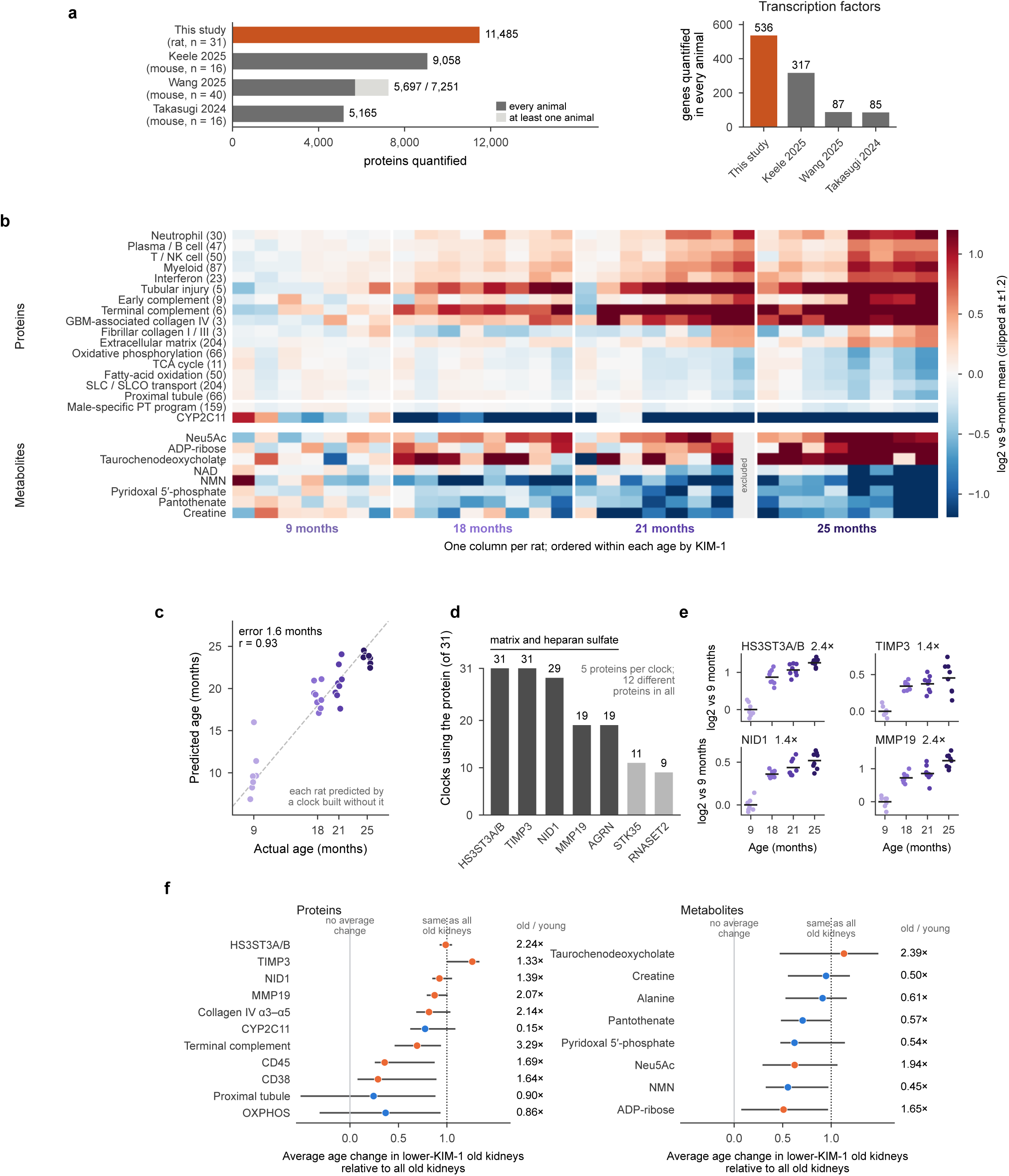
The deep proteome and metabolome of the aging kidney, and a protein clock. (A) Left, proteins quantified in every animal in this study (rat, n = 31) and in three mouse kidney aging proteomes (Keele et al., 2025; Wang et al., 2025; Takasugi et al., 2024); light gray, proteins seen in at least one animal. Right, transcription factors (Lambert et al., 2018) quantified in every animal. (B) Protein programs, CYP2C11 and selected metabolites in each kidney, relative to the 9-month mean. Each column is one rat, ordered within each age by KlM-1. Program rows are the mean of their member proteins (number in parentheses); CYP2C11 is measured from its gene-unique peptides. One 21-month rat was excluded from metabolomics (gray). (C) Protein clock. Each rat’s age was predicted by a clock built without that rat, which chose five proteins from the whole proteome using only the other rats (Methods). Mean absolute error, 1.6 months, compared with 4.7 months for guessing the average age; holding out one or two rats of each age at a time gave similar errors. Dashed line, perfect prediction. (D) Number of the 31 clocks that used each protein (proteins used by at least five clocks). Dark bars, matrix and heparan-sulfate proteins. HS3ST3A/B is measured from peptides shared by HS3ST3A1 and HS3ST3B1. (E) Four clock proteins by age, relative to the 9-month mean; numbers are fold changes at 25 versus 9 months. Points are rats; lines are age-group means. Among rats of the same age, MMP19 correlates with KlM-1 (r = 0.71), TlMP3 correlates inversely (r = –0.58) and HS3ST3A/B does not (r = –0.17). (F) Average age change in old kidneys with lower KlM-1, relative to all old kidneys. Rats aged 21 and 25 months were split within each age at the median KlM-1 level. A value of 1 means the lower-KlM-1 half changed as much as all old kidneys on average; 0 means no average change in that half. Collagen lV α3–α5, terminal complement and CYP2C11 are the signals shown in B. Bars, 95% bootstrap intervals; orange, rises with age; blue, falls. Right, fold change in 21-and 25-month kidneys relative to 9-month kidneys. n = 7, 8, 8 and 8 rats at 9, 18, 21 and 25 months for proteins and 8, 8, 7 and 8 for metabolites.

On average, protein programs changed in consistent directions with age, although individual kidneys varied (**Figure 4B**). Proteins of immune cells and interferon responses, tubular injury, complement and the basement-membrane collagen lV network rose, while proteins of oxidative metabolism, fatty-acid oxidation, solute transport and mature proximal-tubule identity fell (**Supplementary Figure 3**). The male-specific proximal-tubule enzyme CYP2C11 was strongly reduced in most kidneys by 18 months. The metabolome changed in parallel: the NAD^+^ precursor NMN, the cofactor pyridoxal 5’-phosphate, the vitamin pantothenate, and creatine all fell, and NAD^+^ itself fell more modestly, whereas ADP-ribose, the sialic acid Neu5Ac and conjugated bile acids rose. These changes were selective, involving particular cofactor, amino-acid and bile-acid pools rather than the metabolome as a whole (**Supplementary Figure 4A,B**). ln the older age groups, kidneys with more of the injury marker KlM-1 tended to show the immune, injury and metabolic changes most strongly, so the oldest kidneys differed most from one another.

Between 9 and 25 months, 483 proteins (one record per gene) changed significantly (q < 0.01) and by more than 1.5-fold, most of them increases (401 higher, 82 lower; **Supplementary Figure 5A**). Across the proteome, protein changes broadly followed the RNA changes measured in kidneys of 54 other rats from the same colony (Shavlakadze et al., 2026; **Supplementary Figures 5B and 6**). Extracellular proteins were the clear exceptions: collagen lV, complement components and uromodulin rose far more as protein than their RNA predicted, consistent with altered retention or turnover in the tissue, although changes in synthesis cannot be excluded (**Supplementary Figure 5B**; **Supplementary Figure 7A–F**). Among basement-membrane components, collagen lV rose most, particularly the α3 and α5 chains, which with α4 form the glomerular basement membrane network, while the fibrillar collagens changed little (**Supplementary Figure 7G,H**). The fall in oxidative metabolism and transport accompanied the loss of mature proximal-tubule identity, both across ages and among kidneys of the same age, whereas glycolytic, ribosomal and proteasomal proteins did not decline (**Supplementary Figure 8**). The metabolic decline therefore tracks the loss of differentiated proximal tubule rather than a general loss of cellular machinery.

Because these changes were so consistent, we asked whether a kidney’s age could be read from its proteome alone. We built a protein clock with a regularized regression (elastic net; Zou & Hastie, 2005) and tested it by leave-one-animal-out prediction: each rat’s age was predicted by a clock fit only to the other rats, with its own independent selection of five proteins from the whole proteome (Methods). Predicted age differed from true age by about a month and a half on average, compared with almost five months from simply guessing the average age (**Figure 4C**). Holding out one or two rats of each age at a time gave similar errors (Methods). The clocks were not arbitrary fits: although a separate clock was built for each animal, they repeatedly chose the same few proteins, and two of them, the heparan-sulfate sulfotransferase HS3ST3A/B and the matrix-bound metalloproteinase inhibitor TlMP3 (Yu et al., 2000), were chosen by every clock (**Figure 4D**). Most of the recurring proteins act on the extracellular matrix and its heparan sulfate, among them nidogen-1, which links laminin to collagen lV, and agrin, the main heparan-sulfate proteoglycan of the glomerular basement membrane. The four most selected clock proteins in **Figure 4E** each rose steadily from 9 to 18 months and on to 25, so that for each of them every rat aged 18 months or older lay above every 9-month rat. The two proteins used by every clock did not behave like injury markers: on average they changed about as much in old kidneys with lower KlM-1 as in old kidneys overall (**Figure 4F**), and TlMP3 was actually lower in the more remodeled kidneys (**Figure 5C**). MMP19, used by most but not all clocks, did track KlM-1 among rats of the same age (**Figure 4E**). The clock therefore appears to rest mainly on a gradual change in extracellular-matrix proteins whose average age change is evident even in old kidneys with lower KlM-1, although several clock proteins also vary with the remodeling axis among rats of the same age (**Figure 5C**).

**Figure 5.**
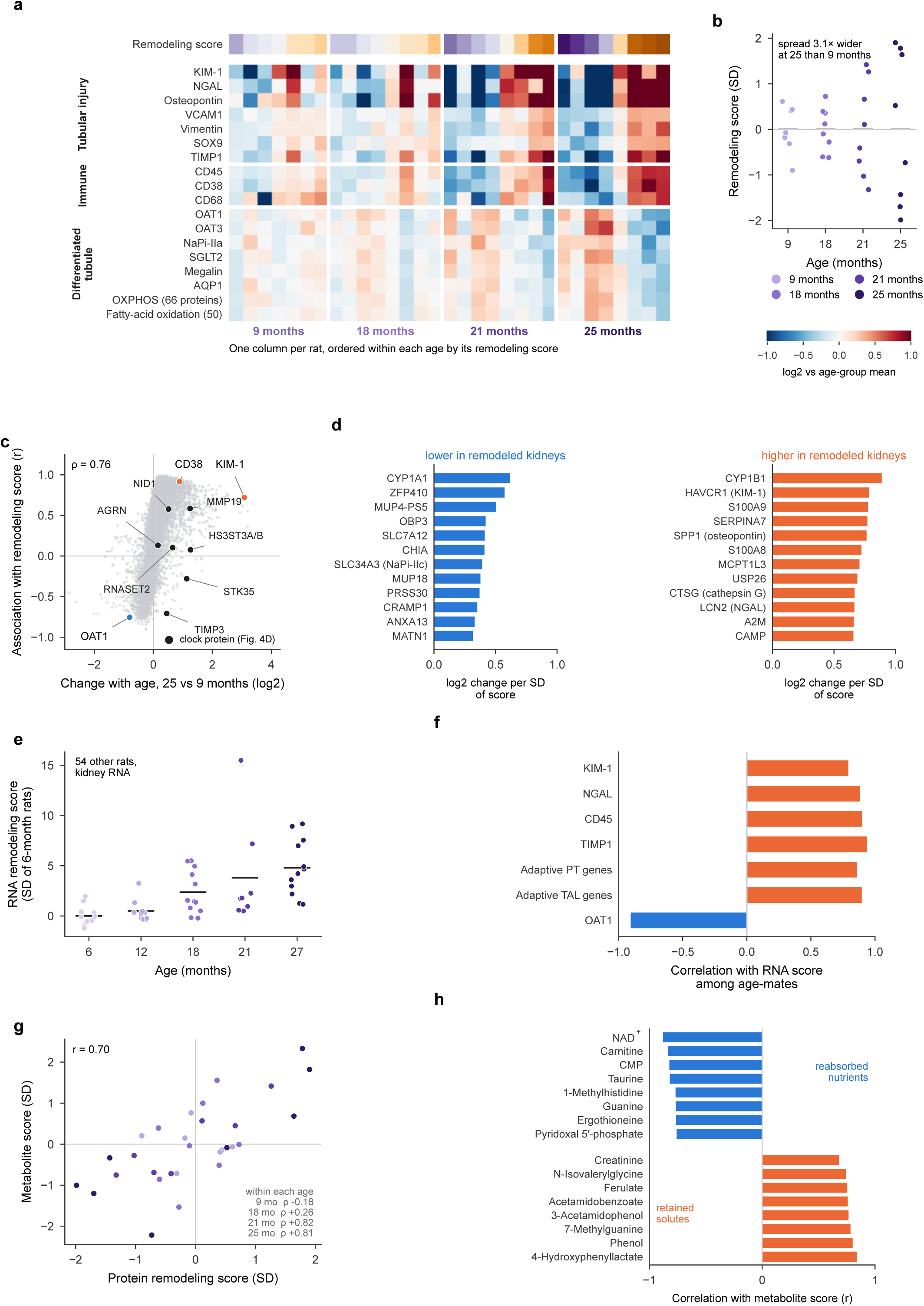
Old kidneys diverge along a dominant injury-remodeling axis seen in protein, RNA and metabolites. (A) Marker proteins in each kidney relative to its own age group (columns are rats, ordered within each age by remodeling score, top strip). The remodeling score is the first principal component of all 11,485 proteins after removing each age group’s mean; higher scores mean more CD45. (B) Remodeling score by age. The spread among rats of the same age is three times larger at 25 than at 9 months. (C) For each protein, its change with age (25 versus 9 months) against its correlation with the remodeling score among rats of the same age (Spearman p = 0.76 across proteins); black, the proteins used by the age clocks (**Figure 4D**). (D) Proteins that change most along the remodeling score (change per SD of score, among named proteins correlated with the score at |r| ≥ 0.5). (E) The same protein pattern applied to kidney RNA from 54 other rats of the same colony (Shavlakadze et al., 2026), relative to 6-month rats. Points are rats; lines are age-group means. (F) Among rats of the same age, correlation of the RNA score with injury, immune and tubule transcripts and with injured-state ("adaptive") proximal-tubule and thick-ascending-limb gene sets (Lake et al., 2023). Each gene or gene set was left out of the score before testing. (G) The metabolome’s own main axis of variation among rats of the same age, computed without the proteins, against the protein remodeling score (30 rats with both measurements; r = 0.70, within-age permutation *P* = 0.001). The two agree in old (21 and 25 months) but not young kidneys. (H) Metabolites most strongly correlated with the metabolite axis, among those annotated at the MS2 level. Proteomics, n = 7, 8, 8 and 8 rats at 9, 18, 21 and 25 months; RNA (E, F), 54 rats; metabolite axis (G, H), 30 rats.

**Figure 6.**
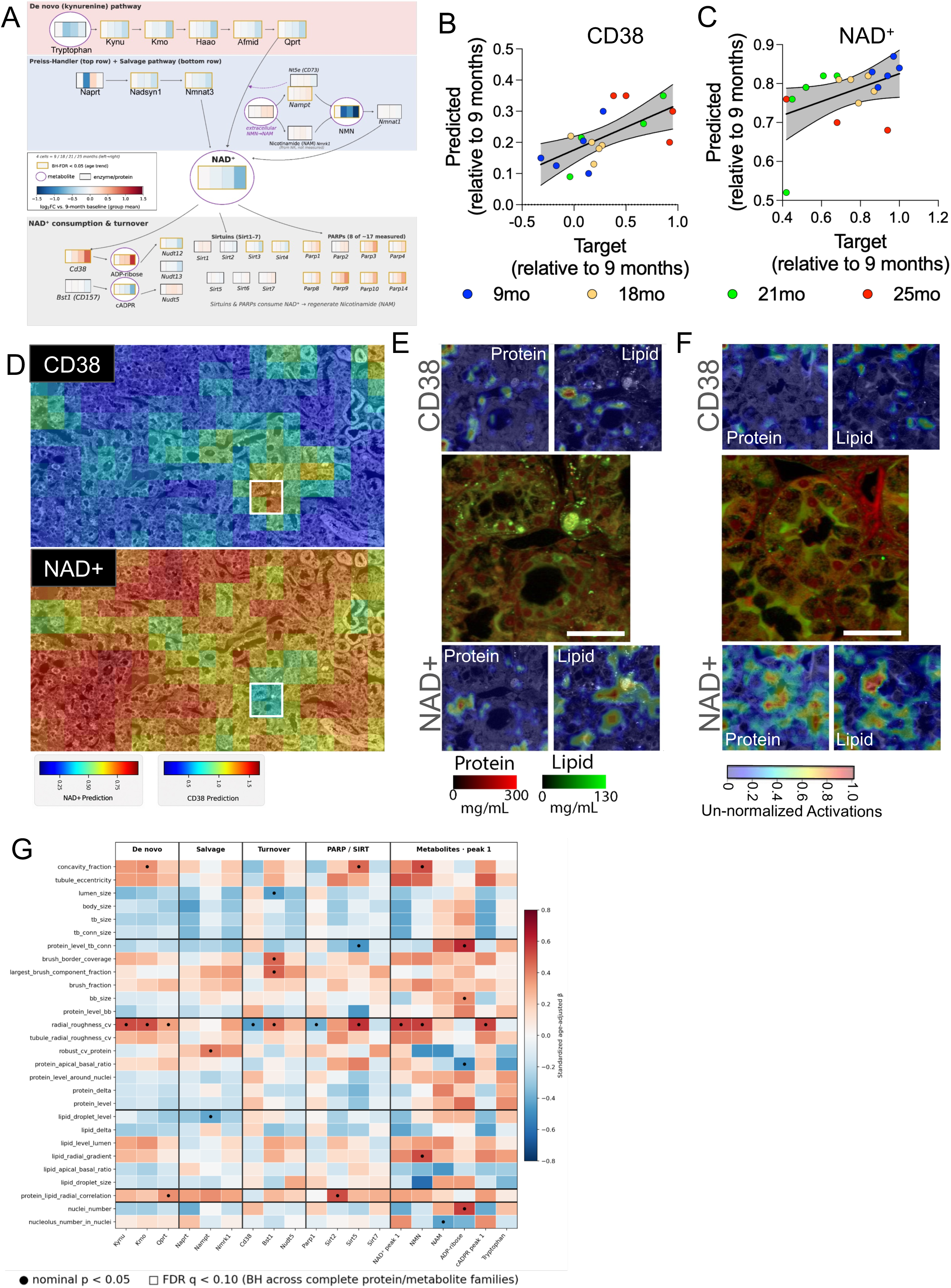
Exploratory links between NoRI morphology and the age-related decline of NAD^+^. (A) NAD^+^ synthesis, consumption and regulation. Each measured protein (black outline) and metabolite (purple outline) is divided into four sections showing its mean log_2_ change at each age relative to 9 months. Orange outlines mark molecules whose abundance correlated with age (Spearman; Benjamini–Hochberg q < 0.05 across all genes or all metabolites; Methods). (B, C) CD38 (B; proteomics) and NAD^+^ (C; metabolomics) abundance predicted from NoRl images of validation animals, against measured abundance, relative to 9 months. Each point is one validation animal (n = 18 in each panel); Spearman’s p is given in the text. (D) Whole-section NoRl image of a 21-month kidney with each tile colored by its predicted CD38 (top) and NAD^+^ (bottom) abundance. White boxes mark the tile with the strongest overlap of high CD38 and low NAD^+^, shown in E. (E) That tile (center) with the CNN activation maps for the features that best predict high CD38 (top) and low NAD^+^ (bottom), in the protein and lipid channels. (F) The same display for a 9-month kidney, with features predicting low CD38 and high NAD^+^. (G) Age-adjusted associations between NoRl features and the abundance of proteins and metabolites of NAD^+^ synthesis (de novo: Kynu, Kmo, Qprt; salvage: Naprt, Nampt, Nmrk1), turnover (CD38, Bst1, Nudt5), PARP/sirtuin pathways (Parp1, Sirt2, Sirt5, Sirt7) and NAD^+^-related metabolites. Colors are standardized regression coefficients (red, positive; blue, negative); black circles, nominal P < 0.05; outlined squares, Benjamini– Hochberg q < 0.10 within the protein or metabolite family. NAD^+^ and cADPR in G are the first of each metabolite’s two chromatographic peaks ("peak 1"). Row labels are the feature names used in the analysis code (defined in **Supplementary Table 1**); those named in the text are radial_roughness_cv (lumen unevenness), concavity_fraction (indentation), tubule_eccentricity (elongation), tb_size (tubule size), tb_conn_size (inter-tubule, interstitial area), protein_level_tb_conn (interstitial protein concentration), lipid_droplet_level (lipid droplet concentration) and protein_apical_basal_ratio (apical-to-basal protein ratio). n = 27– 31 rats per cell, depending on the feature. Concentration and activation scales for E and F are shown below those panels. Scale bars, 40 µm.

KlM-1 is expressed at low levels in the normal kidney but rises sharply in injured proximal tubule cells, in rats and in humans, and is an established marker of proximal-tubule injury (lchimura et al., 1998; Han et al., 2002). Because old kidneys differed widely in KlM-1, we asked which age changes were already evident in old kidneys with lower KlM-1. Within each of the two oldest age groups, we split kidneys at the median KlM-1 level and compared the average age change in the lower-KlM-1 half with that in all old kidneys (**Figure 4F**). On average, the clock and basement-membrane proteins changed about as much in lower-KlM-1 old kidneys as in old kidneys overall, as did creatine, alanine and the bile acid taurochenodeoxycholate. Much of the fall in CYP2C11 and pantothenate and of the rise in complement was also evident in the lower-KlM-1 half. ln contrast, the rise in immune proteins, including CD38, and the loss of proximal-tubule identity and oxidative metabolism were concentrated in the higher-KlM-1 kidneys, although these estimates were less certain. The NAD^+^-related metabolites were intermediate. Aging in the rat kidney therefore appears to combine a broad shift, led by the basement membrane, with KlM-1-associated changes that some old kidneys carry much further than others.

### Old kidneys diverge along a dominant injury-remodeling axis

To study the injury-associated changes directly, we compared each kidney with others of the same age, setting aside the average effect of age. A single pattern dominated (**Figure 5A**): kidneys rich in tubular-injury proteins such as KlM-1 were also rich in immune-cell proteins such as CD45 and CD38, and in the same kidneys transporters and other markers of differentiated tubule, along with the machinery of oxidative metabolism, were low. We summarized this pattern as a remodeling score for each kidney. For every protein we subtracted the average of the kidney’s age group, so that only the differences among rats of the same age remained, and took the main axis of variation (the first principal component) of these differences across the whole proteome (**Supplementary Figure 9**; Methods). Kidneys at 9 months varied little on this score, whereas by 25 months they had spread far apart (**Figure 5B**). Proteins that changed most with age tended to be those that separated more-from less-remodeled kidneys of the same age (**Figure 5C**), consistent with much of the average age response reflecting how far individual kidneys have traveled along this axis. Because these measurements come from whole tissue, the axis could reflect how much of each kidney is injured or infiltrated by immune cells, changes within individual cells, or both; bulk measurements cannot separate the two.

The proteins that rose along the axis included those of injured tubules and of innate immune cells, among them neutrophil and mast-cell proteins (**Figure 5D**); those that fell included male-specific urinary proteins and proximal-tubule transporters. This combination of tubular injury, inflammatory infiltrate and loss of differentiated tubule resembles chronic progressive nephropathy, the common age-related kidney disease of male laboratory rats (Hard & Khan, 2004). The most remodeled kidneys may therefore carry a greater burden of these lesions, although we did not grade them histologically.

To test whether this axis was peculiar to our animals or a general feature of the aging rat kidney, we converted the protein pattern into an RNA signature and applied it to kidney RNA from 54 different rats of the same colony, aged 6 to 27 months (Shavlakadze et al., 2026). The RNA score rose with age, and, as with the protein score, old rats spread widely (**Figure 5E**; **Supplementary Figure 10**; Methods). Among rats of the same age, the RNA score correlated with transcripts of tubular injury, immune cells and injured (“adaptive”) tubule cells and was inversely correlated with the proximal-tubule transporter OAT1, even with each of these genes left out of the signature (**Figure 5F**). The remodeling axis is therefore reproducible in aging kidneys of this rat colony, visible in RNA from independent animals.

Finally, we asked whether the metabolome, measured separately in the same kidneys, would find the same axis on its own. We identified the principal direction in which each kidney’s metabolites differ from those of its age-mates, without reference to the proteins. This metabolite axis agreed with the protein remodeling score, most clearly at 21 and 25 months, when kidneys had diverged most (**Figure 5G**; Methods). At one end of the metabolite axis lay nutrients that the proximal tubule reclaims from the filtrate, such as carnitine, taurine and ergothioneine, together with cellular cofactors, of which NAD^+^ was the most strongly associated (**Figure 5H**). At the other end lay solutes that the kidney normally clears into urine, many of them phenolic compounds from the diet and gut microbes, along with creatinine. The more remodeled kidneys thus contain less of what the tubule normally keeps and more of what it normally excretes, consistent with declining tubular function or reduced clearance. The protein and metabolite axes thus place two molecules at opposite ends: CD38, an NAD^+^-consuming enzyme that rises with immune-cell proteins (**Figure 5A**), and NAD^+^ itself (**Figure 5H**). We examine this pair in **Figure 6**.

### Integrating spatial NoRI and bulk tissue proteomics and metabolomics

A primary function of the NORlGAMl pipeline (**Figure 1**) is to integrate molecular and spatial data to gain deeper insights into the spatial relationships of the circuitry of tissue physiology. NORlGAMl proceeds in three stages: (1) determining molecular (protein/metabolite)–age associations; (2) determining molecular-feature associations; and (3) for candidates identified in steps 1 and 2, we trained neural networks to find spatial dependencies between NoRl features and protein or metabolite abundance within age groups.

We selected the NAD^+^-CD38 axis identified in **Figures 4 and 5** to demonstrate the NORlGAMl integration pipeline, because of its established roles in aging and kidney disease (Camacho-Pereira et al., 2016; Covarrubias et al., 2021) and its prominence in our proteomic and metabolomic analyses. We expanded the axis to incorporate NAD^+^-synthesis arms—*de novo*, Preiss-Handler, and salvage pathways—as well as the consumers and regulators of NAD^+^ to create a global view of NAD^+^ homeostasis and the consequences of its decline in the aging kidney. **Figure 6A** presents a schematic overview of the metabolites and proteins measured in our dataset that are involved in NAD^+^ regulation. The group-mean change in metabolite/protein abundance from 9–25 months is also displayed. This analysis shows that enzymes of de novo NAD^+^ synthesis decline with age, whereas the salvage enzyme NAMPT rises slightly, and that the NAD^+^ consumer CD38 and ADP-ribose, a by-product of NAD^+^ consumption, become more abundant. Among other NAD^+^ consumers, PARPs rose with age and the mitochondrial sirtuin SlRT3 fell, consistent with the metabolic and cell-stress changes that accompany NAD^+^ decline in aging (**Figure 6A**). Thus, aging was associated with coordinated changes in proteins involved in NAD^+^ biosynthesis and consumption, alongside reduced NAD^+^ abundance and increased CD38 abundance, consistent with remodeling of NAD^+^ homeostasis. We therefore integrated these proteins and metabolites with NoRl images from the same animals.

NORlGAMl’s image-based models (a CNN combined with multiple-instance learning; Methods) predict animal-level molecular abundance from NoRl image tiles, as outlined in **Figure 1**. Across the validation animals (three to five per age; 18 animals in each panel), predicted abundance correlated with measured abundance for CD38 (Spearman p = 0.65, *P =* 0.004; **Figure 6B**) and for NAD^+^ (p = 0.48; **Figure 6C**). These correlations span all four ages and therefore partly reflect the shared change of both measurements with age.

We then identified NoRl image features associated with higher CD38 and lower NAD^+^ abundance measured at the animal level. We then mapped the corresponding model-derived molecular signatures onto large-area NoRl kidney images. Regions predicted to have high CD38 overlapped with regions predicted to have low NAD^+^ (**Figure 6D**). These spatial predictions identify candidate regions of molecular alteration rather than directly measured molecular distributions. We extracted the tile with the strongest overlap and used CNN layer 3 activation maps to show the subcellular features that most strongly predict CD38 and NAD^+^ abundance (**Figure 6E**). For comparison, a tile from a 9-month kidney with low predicted CD38 and high predicted NAD^+^ is shown in **Figure 6F**.

We next tested the association of each protein and metabolite of NAD^+^ metabolism with each NoRl feature, adjusting for age (**Figure 6G**). These associations are exploratory and are reported at nominal P < 0.05. The clearest pattern involved the unevenness of the tubule lumen (**Supplementary Table 1**), which was positively associated with the *de novo* NAD^+^-synthesis enzymes Kynu, Kmo and Qprt and with NAD^+^ and NMN, and negatively associated with CD38. Other lumen-shape features (indentation and elongation) tended to relate positively to the same enzymes and to NAD^+^ and NMN, whereas inter-tubule (interstitial) area, lumen size and tubule size tended to relate negatively to several molecules of NAD^+^ synthesis (Kynu, Kmo, Qprt, Naprt, Nmrk1) and turnover (Nudt5). These exploratory analyses also found interstitial protein concentration positively related to ADP-ribose, a product of NAD^+^ consumption by CD38; brush-border coverage positively related to Bst1, an NAD^+^-consuming enzyme; and lipid droplet concentration and the apical-to-basal protein ratio negatively related to Nampt and ADP-ribose, respectively (**Figure 6G**). These associations identify candidates for validation by antibody staining and spatial metabolite measurements on NoRl images of the same samples. Collectively, these exploratory results suggest that NAD^+^-related metabolism may be linked to several aspects of tubular architecture, beyond overall changes in tissue protein or lipid concentration. Some of these features, such as interstitial protein and lipid droplet concentration, can only be measured with quantitative imaging such as NoRl.

To test NORlGAMl in a second setting, we selected kidney injury molecule-1 (KlM-1), an established marker of tubular injury. As in **Figure 6**, we mapped the NoRl features most predictive of KlM-1 abundance. To capture within-age variability, we selected two 21-month kidneys, one with low (**Figure 7A**, top) and one with high (**Figure 7A**, bottom) KlM-1 abundance by proteomics. Predicted KlM-1 abundance tracked measured abundance across the 31 animals plotted (Spearman p = 0.70; **Figure 7B**); as for CD38 and NAD^+^, this correlation spans all four ages. To show tubule subcellular morphology and protein and lipid features that may relate to KlM-1 abundance, we show two NoRl image tiles from 21-month kidneys and their KlM-1 activation maps, representing structures predictive of low (**Figure 7C**) and high (**Figure 7D**) KlM-1 abundance. ln these examples, tubules of normal, healthy appearance were associated with low predicted KlM-1 (**Figure 7C**), whereas regions containing lipid droplets or debris, fragmented tubules and protein plugs were associated with high predicted KlM-1 (**Figure 7D**).

**Figure 7.**
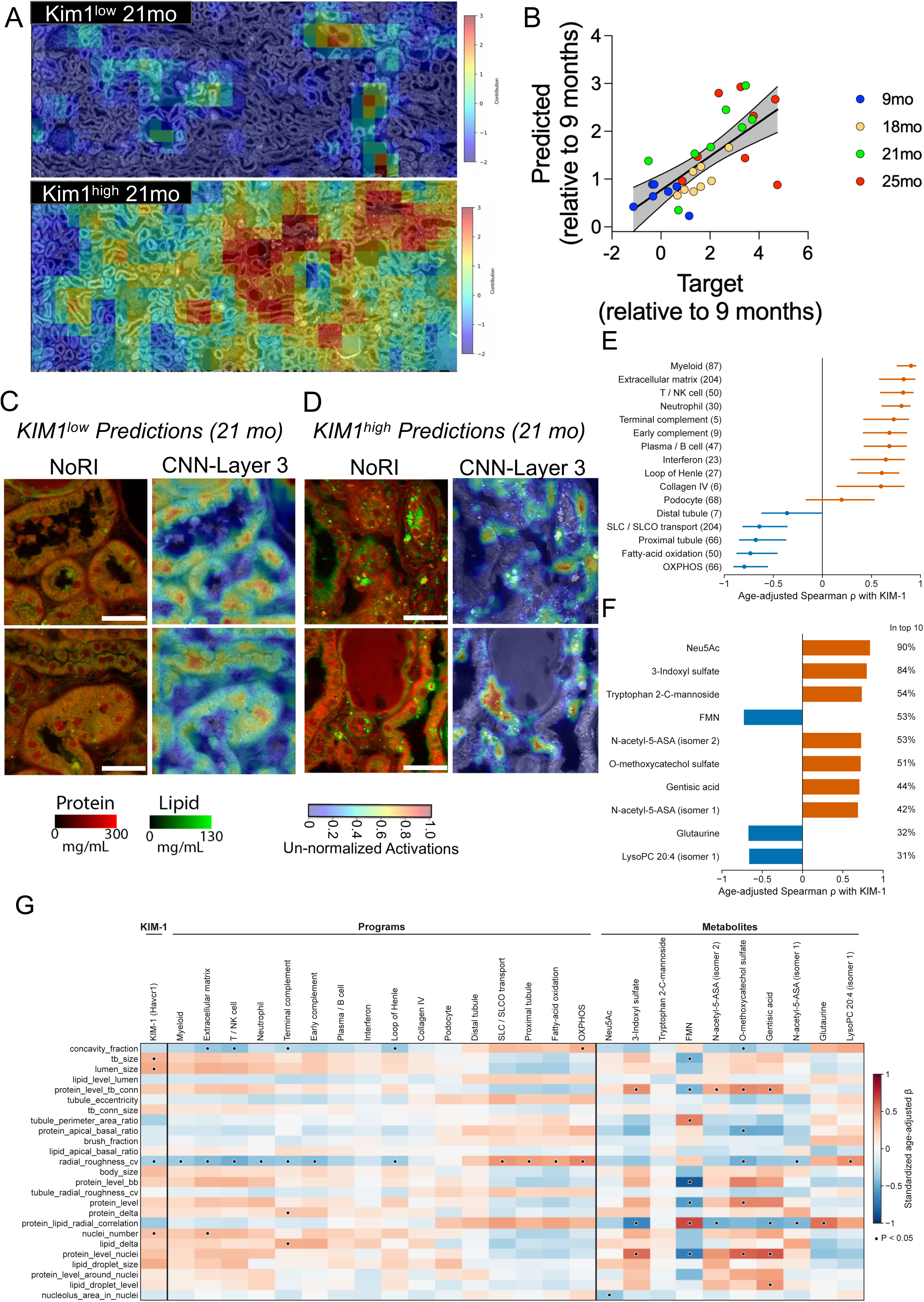
Exploratory links between NoRI morphology and KIM-1 in the aging kidney. (A) Whole-section NoRl images of two 21-month kidneys with low (top) and high (bottom) KlM-1 abundance by proteomics, with each tile colored by its contribution to the predicted KlM-1 abundance. (B) KlM-1 abundance predicted from NoRl images against measured abundance, relative to 9 months; each point is one animal (n = 31). (C, D) Example tiles from 21-month kidneys with CNN activation maps (protein and lipid merged) for the features that best predict low (C) and high (D) KlM-1. (E) Protein programs that track KlM-1 among rats of the same age: age-adjusted Spearman’s p between KlM-1 (HAVCR1) and each program score (number of member proteins in parentheses; KlM-1 is excluded from every program; the terminal-complement program here lacks C8G, which the six-protein score in Figure 4 includes; n = 31). Lines, 95% intervals from 2,000 resamples of rats within age groups; orange, positive; blue, negative. Programs are shown instead of single proteins because 190 proteins correlated with KlM-1 at p ≥ 0.85, so a ranking of individual proteins is unstable. (F) The ten metabolites most strongly correlated with KlM-1 after adjusting for age (n = 30 rats with both measurements); percentages show how often each ranked among the ten strongest in 1,000 resamples of rats within age groups. (G) Age-adjusted associations between NoRl features and KlM-1, the programs in E and the metabolites in F (row labels as in Figure 6G; n = 27– 31 rats per cell, depending on the feature). Colors are standardized regression coefficients (red, positive; blue, negative); black dots, nominal P < 0.05 (exploratory). Concentration and activation scales for C and D are shown below those panels. Scale bars, 40 µm.

Finally, we asked which proteins and metabolites tracked KlM-1 among rats of the same age. Because many single proteins correlated almost equally strongly with KlM-1 (190 at p ≥ 0.85), a ranked list of individual proteins was unstable, so we summarized proteins as predefined programs. KlM-1 tracked immune-cell, extracellular-matrix and complement programs (for example, myeloid p = 0.90) and opposed proximal-tubule, transport, fatty-acid-oxidation and oxidative-phosphorylation programs (p = –0.64 to –0.80; **Figure 7E**), the same pattern as the remodeling axis (**Figure 5A**). Among metabolites, the sialic acid Neu5Ac and the uremic toxin 3-indoxyl sulfate tracked KlM-1 most consistently, whereas FMN was inversely related (**Figure 7F**). We then tested KlM-1, these programs and these metabolites against the NoRl features, adjusting for age (**Figure 7G**). KlM-1 and most immune, matrix and complement programs were negatively associated with lumen unevenness, whereas the proximal-tubule, transport and oxidative programs were positively associated with it, as were NAD^+^ (**Figure 6G**) and LysoPC 20:4. Gentisic acid, an antioxidant reported to protect against renal fibrosis (Wang et al., 2026), tracked KlM-1 and was positively associated with lipid droplet concentration. These associations are exploratory.

These exploratory analyses show that NORlGAMl can relate the morphological and protein– lipid features of kidney aging measured by NoRl to molecular changes in the same animals, providing a foundation for hypothesis generation in aging biology.

## Discussion

This study introduces NORIGAMI, a quantitative framework that links calibrated tissue chemistry and architecture to the molecular biology of renal aging. NoRI identified 41 of 153 morphological and compositional features as significantly age-associated, including anatomically heterogeneous, non-linear lipid remodeling. Deep proteomics (11,485 protein records) and metabolomics (269 metabolites) revealed declining proximal tubule identity and oxidative metabolism, rising injury-associated and inflammatory programs, selective extracellular-matrix remodeling and altered NAD^+^ homeostasis. NORIGAMI predicted the chronological age of held-out rats from NoRI images, with models trained on whole-tissue fields outperforming those trained on isolated tubules, and connected molecular abundance to quantitative tissue features. Two entry points, the NAD^+^ network and the a priori injury marker KIM-1, which lie on the same injury-remodeling axis (**Figure 5**), converged, in exploratory analyses, on shared candidate epithelial phenotypes, suggesting that metabolic decline, lipid handling and injury are linked aspects of renal aging that may be detectable directly in tissue.

A major advantage of NoRI is its ability to quantify tissue protein and lipid concentrations in absolute units (mg/mL) while preserving much of the native architecture. Routine histological processing can extract lipids, distort tissue dimensions and introduce staining variability, leaving lipid-rich structures as apparently empty compartments rather than chemically defined features. This is particularly problematic when interpreting lipid-associated changes in metabolism, inflammation and tissue aging. Unlike conventional stimulated Raman scattering (SRS), whose signal intensities are affected by tissue scattering and attenuation, NoRI uses spectral decomposition to separate protein, lipid and water signals, then normalizes each pixel on the assumption that the volume fractions of these three components sum to unity, yielding absolute concentrations. Because samples need only fixation and sectioning, without dehydration, clearing or staining, these measurements retain compositional information often removed by histology. Daily reference calibration places measurements on a common quantitative scale across imaging sessions, which was particularly important here because acquisition extended over approximately 18 months (Oh et al., 2022). In our initial renal application, quantitative lipid information improved acute kidney injury classification and revealed changes in lipid droplets and brush-border lipid concentration that would be difficult to characterize from morphology alone (Trim, Oh et al., 2026, in press). NORIGAMI extends this foundation by combining quantitative imaging and predictive modeling with deep proteomics and metabolomics to identify age-associated tissue phenotypes and their molecular correlates.

These findings complement advances in computational histopathology. Abila et al. (2026) predicted age from conventional human kidney histology, with validation mean absolute errors of 4.47 years in cortex and 6.08 years in medulla; differences in species, age ranges and validation procedures preclude direct performance comparisons. Unlike stain-dependent images, NoRI provides calibrated protein and lipid concentrations, allowing models to exploit both tissue architecture and quantitative composition, with reduced susceptibility to staining variability that can impair model generalization (Tellez et al., 2019; Lin et al., 2025). While molecular abundance has previously been inferred from histology using spatial transcriptomics or bulk measurements (He et al., 2020; Schmauch et al., 2020), NORIGAMI links molecular predictions to interpretable biochemical and structural features. Nevertheless, image-derived molecular predictions represent associations rather than direct localization of molecules; validation using antibody and in situ measurements remains necessary.

Our renal proteomic dataset quantified 11,485 protein records, representing 11,040 genes, consistently across 31 animals. This coverage exceeded that of comparable filtered whole-kidney aging datasets (Keele et al., 2025; Wang et al., 2025; Takasugi et al., 2024). The additional depth enabled quantification of 536 transcription factors in every animal (**Figure 4A**). Agreement with the rat data from an independent multi-tissue, multi-timepoint transcriptomic atlas of aging (Shavlakadze et al., 2026) supported major biological trends while revealing differences between RNA and protein responses (**Supplementary Figure 5**). This illustrates how deeper profiling can reveal coordinated regulatory and structural processes, while providing the breadth required to relate molecular abundance to quantitative tissue phenotypes with NORIGAMI.

A central biological finding was the coordinated decline in mature proximal tubule identity and oxidative metabolism. Oxidative phosphorylation, fatty-acid oxidation, solute transport and particularly S1-associated proximal tubule markers decreased with age, without comparable declines in glycolytic, ribosomal or proteasomal programs. Mature epithelial identity and oxidative-metabolic scores remained strongly associated among kidneys of the same age (**Supplementary Figure 8C**), suggesting linked variation even among similarly aged kidneys rather than two changes explained solely by chronological age. Injury-associated loss of differentiated tubular functions has also been observed in human kidney studies (Lake et al., 2023), although bulk proteomics cannot distinguish altered cell proportions from cell-intrinsic changes. Concurrent alterations in tubular lipid concentrations and organization provide a complementary structural phenotype. Determining whether these metabolic and morphological changes occur within the same tubular populations, rather than reflecting tissue-wide compositional shifts, requires spatial validation.

Extracellular-matrix remodeling was also prominent, including increased collagen IV, largest for the α3 and α5 chains of the glomerular basement-membrane α3α4α5 network, with smaller increases in laminins and nidogen-1. This differs from reports of declining basement-membrane constituents in aging mouse and human kidney (Randles et al., 2021) but is consistent with markedly prolonged collagen IV half-lives in aged mouse kidney (Preston et al., 2026), which would favor accumulation through reduced degradation rather than increased synthesis. Age-associated structural alterations have also been described in human renal basement-membrane proteins (Eckersley et al., 2023). Divergence of the glomerular α3α4α5(lV) network from published basement-membrane trends may therefore reflect compartment-specific turnover or protein recovery. Increased collagen abundance alone does not establish fibrosis but illustrates why matrix aging should not be treated as a uniform process.

Comparing kidneys of the same age revealed a dominant injury-remodeling axis, seen separately in the proteome, in kidney RNA from other rats and in the metabolome, linking tubular injury and immune proteins with reduced proximal tubule identity, oxidative metabolism, and lower NAD^+^ and related cofactors (**Figure 5**). By contrast, the matrix proteins at the core of the protein aging clock changed, on average, about as much in old kidneys with lower KIM-1 as in old kidneys overall (**Figure 4C–F**). Together, these findings suggest that renal aging combines a broad matrix shift, evident on average even in old kidneys with lower KIM-1, with metabolic decline and injury-associated responses that vary widely between age-matched animals. The latter points to biological heterogeneity that may be obscured by analyses focused exclusively on group-average abundance.

The NAD^+^–CD38 axis illustrates how NORIGAMI relates molecular changes to morphology. NAD^+^ supports redox metabolism and stress responses, while CD38 can contribute to age-related NAD^+^ depletion (Camacho-Pereira et al., 2016; Covarrubias et al., 2020); impaired NAD^+^ biosynthesis has also been implicated in kidney injury (Poyan Mehr et al., 2018). We observed declining NAD^+^ and related metabolites alongside increasing CD38. This is consistent with disrupted NAD^+^ homeostasis, although the data cannot determine whether changes in synthesis, consumption or cellular composition dominate. NORIGAMI’s image models predicted their measured abundances across animals of different ages, and exploratory age-adjusted associations related the broader NAD^+^ network to tubular geometry, brush-border organization and epithelial protein–lipid compartmentalization. Prediction maps highlighted overlapping regions associated with higher CD38 and lower NAD^+^, indicating that related image features inform the two predictions, although these maps do not constitute direct molecular localization. These converging observations motivate spatial validation of NAD^+^-associated epithelial phenotypes.

We selected KIM-1 a priori because of its established association with acute kidney injury, to which the aging kidney is increasingly susceptible. Beyond marking injured proximal tubules, KIM-1 is a phosphatidylserine receptor that confers a phagocytic phenotype on tubular epithelial cells, mediating uptake of apoptotic bodies and oxidized low-density lipoprotein (Ichimura et al., 2008; Mori et al., 2021). It also facilitates endocytic uptake of palmitate-bound albumin, promoting tubular stress, inflammation and fibrosis in experimental diabetic kidney disease (Mori et al., 2021). In our data, KIM-1 and the protein programs that tracked it were associated, in exploratory analyses, with lumen and tubule shape and nuclear number, and the KIM-1-associated metabolite gentisic acid with lipid droplet concentration (**Figure 7G**). KIM-1 itself was not clearly associated with lipid droplet size or concentration. Whether KIM-1 contributes to epithelial lipid uptake during natural aging remains untested, although such a role has been shown in mouse models of diabetic kidney disease (Mori et al., 2021).

Anchoring analyses on KIM-1 also led back to NAD^+^ homeostasis. The redox cofactor flavin mononucleotide (FMN) was among the metabolites most inversely related to KIM-1 within age (**Figure 7F**), and NAD^+^ and KIM-1 showed opposite associations with lumen unevenness (**Figures 6G and 7G**), as expected for two molecules at opposite ends of the same remodeling axis. CD38 rose together with immune-cell proteins (**Figure 5A**), consistent with CD38-expressing immune cells such as the CD38^+^ macrophages described in aging tissues (Covarrubias et al., 2020), which could link NAD^+^ decline to the inflammatory programs observed here. The cyclic ADP-ribose signal tracked NAD^+^ closely and may arise partly from NAD^+^ itself during measurement, so it cannot serve as a readout of CD38 activity. Gentisic acid tracked KIM-1 within age and was associated with higher lipid-droplet concentration (**Figure 7F,G**). Although it attenuates renal fibrosis after unilateral ureteral obstruction by suppressing LDHA-associated glycolysis and H3K18 lactylation (Wang et al., 2026), in rodents it may derive partly from dietary plant phenolics and their metabolism, so its renal abundance may also reflect intake or reduced clearance in older kidneys. These observations suggest shared structural and compositional correlates of epithelial stress and altered NAD^+^ homeostasis. However, neither glycolytic flux nor histone lactylation was measured here, and correlations among measurements from the same animals do not constitute independent biological validation.

## Limitations

This study has several limitations. It is cross-sectional, used only male rats of a single strain, and included seven or eight animals per age group, limiting assessment of biological variability and sex-specific aging. Each age group was collected on a single day, so age cannot be separated from collection-day effects. Models were evaluated with animal-level separation of training and validation data, but checkpoints were chosen on the validation animals, so the reported errors are validation estimates; independent cohorts and other species are needed to establish generalizability and conservation of age-associated features. Proteomic and metabolomic measurements were obtained from contralateral kidneys as bulk averages; they cannot resolve cell-type composition, and image-derived molecular predictions and prediction maps therefore represent associations, partly shared through age, rather than molecular localization. Histopathology was not graded, so the remodeling axis could not be related directly to chronic progressive nephropathy lesions. Antibody-based and spatial metabolite measurements on imaged sections are needed to validate localization, and perturbation experiments to test causality. Given the prominence of lipid phenotypes, the absence of lipidomics is a key omission. Finally, measures of renal function (blood and urine markers) and of organismal aging (e.g., DNA-methylation clocks, metabolic flexibility and immune dysregulation) would anchor tissue phenotypes to physiology.

## Conclusion

NORIGAMI therefore connects quantitative tissue architecture and composition with molecular changes during renal aging. The convergence of declining proximal tubule metabolic identity, heterogeneous lipid remodeling, altered NAD^+^ homeostasis and KIM-1-associated injury responses suggests that these processes represent related aspects of age-dependent tissue remodeling. Although their causal relationships and cellular localization require experimental validation, the identification of candidate shared morphological and compositional correlates provides a foundation for investigating how metabolic dysfunction and epithelial injury become integrated within aging kidney tissue. More broadly, combining quantitative biochemical imaging with molecular profiling offers a means to move beyond descriptive histopathology toward understanding the molecular processes underlying tissue aging.

## Methods and Materials

### Animal maintenance

All procedures involving animals were approved by the Institutional Animal Care and Use Committee of Regeneron (NYS AWP-ID# A-125) and New York Medical College (NYS AWP ID# A-122), NY, USA. Male Sprague-Dawley rats were purchased at 3 to 4 weeks of age from Envigo (Indianapolis, USA) and aged at Envigo under SPF conditions. Rats were imported to New York Medical College animal holding facilities and acclimated for at least 2 weeks. Once imported, the rats were housed two per cage at the SPF facility with controlled temperature and light (22°C, 12-hour light/12-hour dark cycle: lights on at 0600 hours/lights off at 1800 hours) and with ad libitum access to food (2014 Teklad Global 14% Protein diet, Envigo) and water.

### Kidney resection and preparation

Tissue collection occurred between June 2024 and July 2025 in batches, and tissues from the same age group (n = 8 per group) were collected on the same day between 10 a.m. and 1 p.m. Prior to kidney collection, rats were deeply anesthetized with 150 mg/kg ketamine (Ketaset; Pfizer, USA) and 10 mg/kg xylazine (Anased; Lloyd Laboratories, USA) followed by transcardiac perfusion (into the aorta via the left ventricle) with 200 mL of filter-sterilized phosphate buffered saline (PBS) containing 20 U/mL heparin (Millipore-Sigma, USA) cooled to 4 °C. Kidneys were resected following perfusion: the left kidney was used for imaging and the right kidney, from the same animal, for mass spectrometry analysis. For mass spectrometry analysis, kidneys were immediately snap frozen in liquid nitrogen following resection and then transported and stored at –80 °C until analysis.

### Sample allocation, exclusions and experimental design

Thirty-two male rats were collected (n = 8 at each of 9, 18, 21 and 25 months). One 9-month animal was excluded from proteomics because of a visibly bloody preparation, leaving 31 proteomes; N151, which had the highest erythrocyte signal of these, was retained (**Supplementary Figure 11B**). A different animal’s metabolome was excluded because of a sample run/injection error, leaving 31 metabolomes. Thirty animals had both datasets and were used for all paired analyses. NoRI data were obtained for all 32 animals.

### Proteomics sample processing and data acquisition

Ages were balanced across TMT series and preparation batches (**Supplementary Figure 11A**); neither was fitted as a model term. Animals are the biological replicates throughout; fractions, peptide forms, sequences and images do not increase biological n.

#### Sample preparation for LC-MS

Lysates were sonicated on ice for 10 cycles of 15 s at 60% amplitude, then clarified by centrifugation at 20,000 × g for 15 min at 4 °C. Disulfide bonds were reduced with 5 mM dithiothreitol (DTT; Oakwood Chemical) for 15 min at 60 °C. Free cysteines were alkylated with 50 mM iodoacetamide (Millipore Sigma) for 1 h in the dark, and the reaction was quenched with 30 mM DTT. Proteins were then precipitated with chloroform/methanol (Wessel & Flügge, 1984). The resulting protein pellet was resuspended at approximately 2 µg/µL in 6 M guanidine hydrochloride (GuHCl) containing 10 mM EPPS, pH 8.5.

For each sample, 50 µg of protein was diluted to 2 M GuHCl with 10 mM EPPS, pH 8.5, and digested with LysC (20 ng/µL; Wako) for 14 h. The digests were further diluted to 0.5 M GuHCl and supplemented with trypsin (10 ng/µL; sequencing grade, Promega) while maintaining LysC at 20 ng/µL, then incubated for 16 h at 37 °C. After the solvent was evaporated under vacuum, peptides were resuspended at approximately 1 µg/µL in 500 mM EPPS, pH 8.0.

To make the bridge samples, equal volumes of all 31 rat tryptic digests were pooled, mixed, and split equally into four tubes. These were labeled with TMTpro-126, TMTpro-135N, TMTpro-127D or TMTpro-135CD. Each 50 µg peptide aliquot was labeled with 20 µL of TMTpro-D or TMTpro-H reagent (Thermo; 20 µg/µL in anhydrous acetonitrile, ACN) for 2 h at room temperature. The reaction was quenched with 10 µL of 5% hydroxylamine (Millipore Sigma).

Labeled samples were pooled and dried under vacuum. The pool was acidified to pH ≤ 1 with TFA and clarified at 20,000 × g for 10 min at 4 °C. It was then desalted on a Sep-Pak cartridge (Waters), which was conditioned with 1 mL methanol and equilibrated with 1 mL of 1% formic acid (FA). After loading, the cartridge was washed twice with 1 mL of 1% FA, and peptides were eluted with 1 mL of 35% ACN/1% FA into a 2 mL Eppendorf tube. The eluate was dried under vacuum and resuspended in 140 µL of 10 mM ammonium bicarbonate, pH 8.0. It was then clarified by ultracentrifugation at 70,000 rpm for 30 min at 4 °C. A 100 µL portion of the supernatant was separated by basic-pH reversed-phase chromatography as described previously (Navarrete-Perea et al., 2018), collecting into a 96-well plate and combining into 23 fractions. Fractions were desalted on StageTips (Rappsilber et al., 2007), and approximately 10 µg of peptides per fraction was analyzed by LC-MS/MS.

#### LC-MS/MS

Data were acquired on a Thermo Orbitrap Eclipse coupled to a Thermo EASY-nLC 1200, operated in data-dependent mode. Each run lasted 7 h, with a gradient from 3% to 18% B. Solvent A was 0.125% FA and 2% DMSO; solvent B was 0.125% FA, 80% ACN and 2% DMSO.

Peptides were separated on in-house packed microcapillary columns (100 µm inner diameter, 360 µm outer diameter). Each column was packed by hand with about 0.5–1 cm of Magic C4 resin followed by 40 cm of 1.9 µm C18-AQ resin (Dr. Maisch GmbH, part no. r119.aq.0001), as described previously (Sonnett et al., 2026). Columns were held at 60 °C in a custom-machined heater. Electrospray ionization used 2.6 kV applied through a PEEK microtee.

Survey scans covered 300–1,100 m/z at 120,000 resolution in the Orbitrap, with an AGC target of 1 × 10^6^, a maximum injection time of 200 ms, a 10 s cycle time and an RF lens setting of 60%. Each run comprised four experiments, one at each FAIMS (Schweppe et al., 2019) compensation voltage (–32, –42, –52 and –62 V). FAlMS was run at standard resolution with a total carrier gas flow of 3.5 L/min. Dynamic exclusion lasted 90 s with a ±10 ppm tolerance.

For MS2, precursors were isolated in the quadrupole with a 0.4 m/z window and fragmented by CID at 35% normalized collision energy. MS2 spectra were recorded in the Orbitrap at 15,000 resolution with a first mass of 110 m/z, an AGC target of 7.5 × 10^4^ and a maximum injection time of 50 ms.

TMTpro 35-plex reporter ions were quantified in an SPS-MS3 scan (McAlister et al., 2014) with five synchronous precursors. Isobaric tag-loss ions were excluded from SPS selection. The MS1 isolation window was 0.7 m/z, and fragmentation was by HCD at 45% normalized collision energy. MS3 spectra were recorded in the Orbitrap at 60,000 resolution with TurboTMT enabled, an AGC target of 2.5 × 10^5^ and a maximum injection time of 118 ms. The low-m/z TMTpro reporter ions were extracted with a 20 ppm tolerance and corrected for isotopic impurities as described previously.

#### Data analysis

Peptide sequences were assigned to MS2 spectra with the Comet search engine (Eng et al., 2013), using a 50 ppm precursor mass tolerance. Variable modifications were methionine oxidation (+15.99491 Da) and asparagine deamidation (+0.98402 Da). Static modifications were carbamidomethylation of cysteine (+57.02146 Da) and TMTpro (+304.207146 Da) on peptide N-termini and lysines. Peptide identifications were filtered to 1% FDR using the target-decoy strategy (Elias & Gygi, 2007), then filtered further to 1% FDR at the protein level with a picked target–decoy approach (Huttlin et al., 2010; Savitski et al., 2015). The eight exports, each filtered by the search pipeline to a 1% protein-level FDR, contained 11,744 rat target and 129 rat decoy protein identifiers, excluding contaminants.

#### Proteome quantification and molecular identity

TMTpro reporter-ion signal-to-noise (S/N) values (Li et al., 2020; Zuniga et al., 2024) were taken from the search-pipeline export, which was already corrected for isotopic impurities. PSMs with summed raw S/N of at least 100 in both the H and the D series were used. Within each series, channels were scaled to equal total S/N. For each PSM, the D channels were then multiplied by the ratio of the mean H bridge (126, 135N) to the mean D bridge (127D, 135CD), placing both series on one scale (Plubell et al., 2017). After removing decoy and contaminant matches and four PSMs with an invalid bridge pair, 411,774 PSMs remained.

PSM S/N was summed for each protein, the bridge channels were removed, and values were log_2_-transformed and centered on each protein’s mean in the seven 9-month animals. Of 11,528 proteins quantified, 11,485 had nonzero signal in all 31 animals (in TMT data a zero is a measured value); these 11,485 were used for all log_2_ analyses, without imputation (**Supplementary Table 2**). Where one value per gene was needed (gene-level counts and RNA comparisons), one designated record represented each gene (11,040 genes), usually the one with the most distinct peptide sequences, then the most PSMs.

Peptide sequences were matched to the target FASTA with isoleucine/leucine equivalence, and a sequence was called gene-unique if it matched only one gene. Six signals without a resolving sequence are reported under group names: HS3ST3A1/HS3ST3B1, renin family, C4/C4A, CYP4A2/CYP4A2L1, PSMB6/PSMB6L1 and MUP18/MUP4L4. Protein values use all peptides assigned to the protein unless a gene-unique reconstruction is stated. The CYP2C11 profile (**Figure 4B**) was rebuilt from its 11 gene-unique sequences, summing linear signal before log transformation.

### Metabolomics

Metabolomics was performed as previously described, with minor modifications (Xiao et al., 2025; Yu et al., 2020; Zhang et al., 2024). Metabolites from tissues were extracted by homogenizing with solvent (90% methanol, 10% water) at 1:4 tissue weight-to-solvent volume. Three 20 s cycles at 6,800 RPM with a 30 s pause between cycles on a Precellys Evolution homogenizer (Bertin Technologies) were used. The samples were precipitated at –80 °C overnight, and the supernatant was collected for subsequent LC-MS analyses after centrifuging at 17,000 × g for 15 min at 4 °C.

LC-MS analyses of metabolite extracts were performed on a Vanquish HPLC System controlled by Chromeleon software coupled to an Exploris 120 mass spectrometer equipped with an HESI ion source controlled by Xcalibur software (Thermo Fisher Scientific). Metabolites were separated on an ACQUITY UPLC HSS T3 column (150 mm × 2.1 mm, particle size 1.8 µm) using mobile phase A/B acetonitrile/water (both with 0.1% formic acid). The column was maintained at 35 °C, flow rate was set to 0.4 mL/min, and an A/B gradient was as follows: being isocratic at 1% B for 5 min, linearly increasing to 99% B over 12.5 min, being isocratic at 99% B for 3.5 min, shifting back to 1% B in 0.1 min, and holding at 1% B until 25 min. For quantification in individual samples, the mass spectrometer was operated in positive/negative switching electrospray ionization (ESI) mode (spray voltage +3.5 kV for ESI+ and -2.5 kV for ESI-) scanning MS1 m/z 70–800, using a resolution setting of 120,000 and standard AGC target. For identification, MS1–MS2 data were acquired for six random samples using either ESI+ or ESI-: for MS1, the Orbitrap was operated at a resolution of 60,000; for MS2, top 4 ions at an isolation window of 1.2 m/z with 10 s exclusion (± 5 ppm) were triggered, Orbitrap resolution was 30,000, and stepped collisions were applied (20%, 40%, 60%, normalized). The following mass spectrometer settings were used for both methods: ion transfer tube temperature 320 °C; vaporizer temperature 350 °C; sheath gas 50 AU; auxiliary gas 10 AU; sweep gas 1 AU; RF Lens 70%. The mass spectrometer was calibrated with FlexMix calibration solution (Thermo Fisher Scientific).

Raw LC–MS data files, including MS1 data from biological samples, MS1–MS2 data acquired for compound identification, and solvent-only blank runs, were processed using Compound Discoverer (Thermo Fisher Scientific, version 3.3) with the default metabolomics workflow, incorporating database searches against mzCloud, Mass Lists, and ChemSpider. Putatively annotated compounds with a peak shape score ≥ 3 in at least 80% of samples within at least one biological group were selected for manual curation. Manual curation involved (i) evaluation of MS1-level evidence, including mass accuracy and isotopic patterns relative to predicted elemental compositions; (ii) comparison of experimental MS2 spectra with reference spectra from authentic standards available in mzCloud; and (iii) assessment of the biological plausibility of the proposed compound annotations. Identification confidence was defined according to Schymanski et al. (2014).

### Proteomic and Metabolomic Analysis Procedures

#### Software and general statistical conventions

Proteomic and metabolomic analyses (**Figures 4, 5 and 7E–G**; **Supplementary Figures 3– 11**) used Python 3.12.7 (NumPy 1.26.4, pandas 2.2.2, SciPy 1.13.1, scikit-learn 1.5.1, statsmodels 0.14.2, matplotlib 3.9.2). Other analyses used R 4.5.3 and Python 3.11.15 (numpy 2.4.6 (Harris et al., 2020), pandas 2.3.3 (McKinney, 2010), scipy 1.17.1 (Virtanen et al., 2020), scikit-learn 1.9.1 (Pedregosa et al., 2011), statsmodels 0.14.6 (Seabold & Perktold, 2010), h5py 3.16.0). Image segmentation and deep-learning analyses additionally used PyTorch (Paszke et al., 2019), Ultralytics YOLOv8 (Jocher et al., 2023), UMAP (umap-learn; McInnes et al., 2018) and a ResNet-50 architecture (He et al., 2016). Unless stated otherwise, multiple-testing correction used the Benjamini-Hochberg (BH) false discovery rate (Benjamini & Hochberg, 1995) within each analysis, and permutation tests were used wherever a method’s null distribution is non-standard.

#### Statistical models and program scores

Protein, program and metabolite effects use animal-level ordinary least squares with categorical age (9 months as reference) and HC3 standard errors (MacKinnon & White, 1985). No TMT-series, preparation-batch or other covariates were fitted. Benjamini–Hochberg correction is applied within the stated family: all 11,485 proteins (gene-level displays use these record-level *q* values), all 269 metabolites, or the defined program or outcome family. Highlighted changes require q < 0.01 and, where stated, at least a 1.5-fold change. NAD^+^ is the sum of its two chromatographic peaks and is reported with its nominal *P* value. Correlations are Pearson unless stated otherwise.

Program scores are means of member log_2_ abundances, with memberships fixed before analysis (member-level data in **Supplementary Figure 3**). They describe relative recovered abundance, not cell fractions or pathway activity. Human kidney cell-type and injury-state markers come from Lake et al. (2023), mapped to rat genes; androgen-supported sets come from Xiong et al. (2023). The erythrocyte-associated (RBC) score is the median across-animal z score of the hemoglobin proteins (taken as one component) and of erythrocyte proteins whose profiles tracked hemoglobin consistently (Pearson r ≥ 0.75 across all animals, ≥ 0.70 without N151, and Spearman p ≥ 0.55). A plasma-associated score summarizes predefined plasma proteins (**Supplementary Figure 11B,C**). Neither score measures perfusion quality directly.

#### Proximal tubule (PT) conditioning and program-score controls

PT conditioning adds an external PT marker score to the age model after removing markers that overlap the target program (66 markers remain for OXPHOS, 64 for FAO and 45 for SLC transport). The remaining-difference percentage is 100 × the 25-versus-9-month contrast with the PT score divided by the original contrast; negative values indicate sign reversal. It does not measure a biological or causal fraction. As a specificity control, 4,000 random sets of 66 proteins were drawn from proteins with 25-versus-9-month effects between –0.6 and –0.05 log_2_, excluding OXPHOS and PT members; the 3,626 sets whose mean age effect was within 0.03 of the PT score’s were each used in place of the PT score (**Supplementary Figure 8F**). Within-age covariance (**Supplementary Figure 8C**) uses deviations from age-group means.

#### Terminal complement and UMOD analyses

The terminal-complement score is the mean of C5, C6, C8A, C8B, C8G and C9. Plasma-relative excess is the within-animal difference between this score and a comparator score: a 16-protein plasma set, albumin alone or C3 (**Supplementary Figure 7C**). UMOD peptides were mapped to the 644-residue rat precursor and grouped as upstream or C-terminal of R591, a boundary indirectly supported by rat urinary peptides (Mary et al., 2017); the one sequence spanning it was excluded. Within each sequence, peptide forms were weighted by PSM count; sequences were then averaged with equal weight within each region, and the regional contrast is upstream minus C-terminal. UMOD regions are shown in **Supplementary Figure 7D–F**.

#### Comparison with published kidney proteomes

Coverage (**Figure 4A**) counts protein records in each released matrix: for this study, proteins quantified in all 31 rats (11,485; 11,528 in at least one rat); for Keele et al. (2025), the authors’ filtered kidney matrix (9,058 records, all complete across 16 mice); for Wang et al. (2025), the pre-imputation kidney matrix (5,697 complete and 7,251 observed records across 40 mice); and for Takasugi et al. (2024), the whole-tissue lysate table (5,165 complete records across 16 mice). Complete means an observed, finite value in every animal before any imputation. Transcription factors (**Figure 4A**) are counted as distinct genes among complete records. Transcription factors are members of the human TF census of Lambert et al. (2018, v1.01), transferred to rat and mouse through MGI homology classes; a rodent gene was counted only if every human member of its homology class is a census TF. Records whose gene attribution was ambiguous were not assigned to a class.

#### Cross-cohort RNA comparisons

The rat RNA cohort comprises 54 male kidneys from the same colony at 6, 12, 18, 21 and 27 months (n = 11, 11, 12, 8 and 12) (Shavlakadze et al., 2026). Deposited TMM-normalized (Robinson & Oshlack, 2010) log_2_-CPM data were matched to the protein gene representatives, giving 9,580 shared expressed genes. RNA 27-versus-6-month effects use the source age-group analysis with Welch’s test (Welch, 1947); protein effects compare 25 with 9 months. Huber regression (Huber, 1964) describes the global protein–RNA relationship, and protein residuals from that fit define protein–RNA departures (**Supplementary Figures 5B and 7A**). For trajectories (**Supplementary Figure 6**), each layer is centered on its own youngest group. The animals differ between cohorts, so these are cohort-level comparisons, not paired protein–RNA regulation.

#### Protein clock

Each rat’s age was predicted by a model built without that rat (leave-one-animal-out). Within each training set, proteins were standardized, the penalty of an elastic net (mixing parameter 0.5; Zou & Hastie, 2005) was chosen by four-fold age-stratified cross-validation, and proteins were ranked by how often the elastic net selected them across 12 age-balanced half-samples (Meinshausen & Bühlmann, 2010). The five most often selected proteins were combined by ordinary least squares. The held-out rat was never used for scaling, tuning or selection, so selection cannot bias the error estimate (Varma & Simon, 2006). Clocks of three to ten proteins gave similar errors. To test larger held-out sets, the whole procedure was repeated with splits that held out one or two rats of each age at a time (ten random repeats each; with only seven 9-month rats, one split per repeat was one rat smaller); across repeats, the median of the mean absolute error was 1.6 and 1.7 months.

#### Age changes in old kidneys with lower KIM-1

Rats aged 21 and 25 months were split within each age at the median KIM-1 (HAVCR1) abundance, with rats at the median assigned to the lower half; these halves were identical to those from a split at the median remodeling score. For each protein, program or metabolite, the ratio (mean of the lower-KIM-1 old rats – 9-month mean) / (mean of all old rats – 9-month mean) compares subgroup-average changes: 1 means the lower-KIM-1 half changed as much as all old rats on average, and 0 means no average change in that half; it does not describe individual kidneys. The collagen lV α3–α5 (COL4A3–COL4A5), terminal complement, program and CYP2C11 signals are those shown in **Figure 4B**. Intervals are the 2.5th–97.5th percentiles of 2,000 bootstrap resamples of rats within age groups, with the KIM-1 median recomputed in each resample (Efron & Tibshirani, 1993).

#### Remodeling score

Each protein’s age-group means were subtracted and its deviations scaled to unit variance. The remodeling score is the first principal component of this 31 × 11,485 matrix, oriented so that higher CD45 (PTPRC) is positive (**Supplementary Figure 9**). In **Figure 5C**, each protein’s association with the score is the correlation of its within-age deviations with the score; in **Figure 5D**, named proteins (uncharacterized loci, immunoglobulin and MHC records excluded) correlated with the score at |r| ≥ 0.5 are ranked by the slope of their deviations on the score.

#### RNA remodeling score

Protein loadings on the remodeling score were mapped to genes by gene name (for a gene with several protein records, the first record in the protein table was used) and restricted to genes measured in the RNA data; the 300 most positive and 300 most negative genes were given equal positive or negative weights, which were applied to standardized kidney RNA from the 54-rat cohort above. Marker associations (**Figure 5F**) are within-age correlations computed with each marker gene or gene set (Lake et al., 2023) left out of the score. Three other weightings gave nearly identical scores (**Supplementary Figure 10**).

#### Metabolite axis

For the 30 rats with both datasets, log_2_ metabolite abundances (cyclic ADP-ribose features, which track the NAD^+^ signal, omitted) were centered on age-group means and scaled to unit variance. The metabolite score is the first principal component of this matrix, computed without the proteome. Its agreement with the protein remodeling score was tested by permuting the protein score within age groups (9,999 permutations). **Figure 5H** shows the metabolites most strongly correlated with the score among those annotated at the MS2 level.

#### Multiple testing and permutation tests

Unless otherwise specified, statistical hypotheses were assessed at the biological-animal level; multiple-testing correction used the Benjamini–Hochberg false discovery rate procedure, applied within the analysis family stated above.

### Normalized Raman imaging (NoRI) of kidney tissue

#### Tissue fixation and slicing

For imaging by NoRI, kidneys were cut into four pieces following the lateral axis and were submerged into 50 mL of 4% paraformaldehyde (Thermo Fisher Scientific; USA) at 4 °C for 24 hours, after which samples were washed with 4 °C PBS and finally submerged in 30 mL 4°C PBS containing 0.02% (w/v) sodium azide (Sigma-Aldrich; USA) for shipment to Boston, MA. NoRI imaging was performed on the mid-pole region of the left kidney. Samples were stored for no longer than two weeks before processing for NoRI imaging.

Whole kidneys were removed from storage and placed into mounting cylinders compatible with the Precisionary ZF-310-0F Compresstome® (Precisionary Instruments LLC.; MA, US) containing 2.5% agarose diluted in distilled water (pre-heated to 55 °C) (EZ Pack Agarose™; Benchmark Scientific, Inc.; NJ, US). All buffers used were filtered through a 0.22 µm filter (Corning®; AZ, US) prior to use. Constructs were then kept at 4 °C for 30 minutes to harden the agarose-sample construct. Samples were next placed into the Compresstome® and cut into 120 µm slices at speed setting 3, oscillation setting 5. lndividual tissue slices were then placed into ∼2 mL chilled PBS (0.02% sodium azide) within a 12-well plate (VWR International, LLC; NJ, US).

#### Immunofluorescence staining and mounting

Where immunofluorescence was performed, following Compresstome slicing, samples were placed into PBS containing 5% species-specific normal serum (Jackson ImmunoResearch Laboratories Inc.; PA, US), selected according to the host species of the secondary antibodies used in each staining panel (donkey or goat serum), and 0.02% sodium azide overnight at 4°C with gentle rocking. Blocking solution was removed and replaced with PBS containing 0.02% sodium azide at room temperature with gentle rocking for 20 minutes, repeated three times. After washing, the wash buffer was replaced with primary antibodies in tandem in PBS containing 0.02% sodium azide for combinations of the following antibodies: Anti-uromodulin (UMOD), clone EPR20071, rabbit monoclonal (Abcam, ab207170; 1:200); Anti-aquaporin-1 (AQP1), rabbit, CoraLite Plus 488-conjugated (Proteintech, CL488-20333; 1:250); Anti-aquaporin-2 (AQP2), goat (Thermo Fisher Scientific, OSA00145W; 1:200); Anti-podoplanin (T1α), purified anti-rat, mouse IgG1 (BioLegend, 250902, E11; 1:250); Anti-KIM-1/TIM-1/HAVCR1, rabbit (R&D Systems, AF1817; 1:200); and Anti-CD38, purified anti-rat, mouse lgG2b, k (BioLegend; 250502, T10; 1:200). Samples were incubated in primary antibody cocktail for 24 hours at room temperature with gentle rocking. Primary antibody cocktail solution was then removed and replaced with PBS containing 0.02% sodium azide for 30 minutes at room temperature with gentle rocking to wash. This wash was repeated 4–6 times. Samples were incubated in secondary antibody cocktail solution for 24 hours at room temperature with gentle rocking, in darkness, in PBS containing 0.02% sodium azide with combinations of the following antibodies: donkey anti-goat IgG (H+L), DyLight 405; goat anti-mouse IgG1, Alexa Fluor 488; and donkey anti-rabbit IgG (H+L), DyLight 594. All secondary antibodies were purchased from Jackson ImmunoResearch Laboratories Inc. and used at 1:500 dilution. Secondary antibody cocktail solution was removed and replaced with PBS containing 0.02% sodium azide for 30 minutes at room temperature, in darkness, with gentle rocking, repeated 4–6 times. For nuclei segmentation training two samples per age group were stained with 9 µg/mL Hoechst 33342 (Thermo Fisher Scientific; USA) for 1 hour at 4 °C with gentle agitation, followed by four 30-minute washes in PBS containing 0.02% sodium azide.

After washing, samples were mounted onto glass slides (Superfrost® Plus VWR MicroSlides; VWR International LLC.; USA) in anti-fade mounting solution containing 0.2% propyl gallate (Sigma-Aldrich; USA) diluted in 0.8% fully deuterated dimethylsulfoxide (DMSO; Cambridge Isotope Laboratories Inc.; USA) in PBS containing 0.02% sodium azide. The deuterated form of DMSO was used so as not to contribute to the NoRI quantification of the protein methyl groups. Number 1.5 micro cover glass slides (VWR International, LLC.; USA) were used to cover samples and slides were sealed using clear nail polish. Samples were stored at 4 °C in darkness in a humidified container until imaging.

#### Fluorescence microscopy and normalized Raman imaging (NoRI)

Microscopy imaging and quantitation was performed on an Olympus FV3000 using FluoView (Olympus; Tokyo, Japan) and MATLAB (MathWorks; US). Immunofluorescence imaging was used for cell-type and tubule identification and for identification of markers of cellular stress. Immunofluorescence imaging was followed by NoRI imaging for protein and lipid quantification. All images were taken using a water immersion 60×/1.2 NA objective lens (Evident Scientific) and an oil immersion 1.4 NA condenser lens, where each pixel was equivalent to 414 nm for both NoRI and immunofluorescence imaging.

Protein and lipid concentrations were calculated from SRS images using NoRI as previously described (Oh et al., 2022). Briefly, SRS images at 2853 cm^-1^, 2935 cm^-1^, and 3420 cm^-1^ bands (corresponding to the methylene- and methyl-groups, and water characteristic vibrational bands, respectively) were acquired from fixed tissue slices using a custom-built spectral-focusing femtosecond SRS microscope. This microscope was constructed using synchronized femtosecond pulse lasers for the Pump and Stokes beams. A pair of dense flint (DF) glass rods chirped the pulses. An electro-optical modulator (EOM) modulated the amplitude of the Stokes beam at 20 MHz. A retro-reflector prism mounted on a motorized delay was adjusted to control the overlap of the Pump and Stokes beams. The Pump and Stokes beams, combined by a dichroic mirror (DM), were focused on the sample by the objective lens. Images were acquired by point-scanning using a pair of galvanometer scan mirrors. After stimulated Raman scattering at the sample plane, the Pump beam was collected by a high numerical aperture condenser lens, selected by a short pass filter (SF), and its intensity was measured by a high-speed photodetector. The three SRS images were spectrally unmixed into protein, lipid, and water components using reference spectra measured from bovine serum albumin solution in water, dioleoyl-phosphocholine solution in per-deuterated methanol, water, and per-deuterated methanol. Unmixed images of protein, lipid, and water components were converted to the absolute concentration by using the sum of the three components as the normalization reference at each pixel.

#### NoRI imaging and image preprocessing

Imaging parameters and preprocessing were kept constant across all regions (Oh et al., 2022). For analysis, two-dimensional NoRI images were generated by applying maximum-intensity projection. Protein and lipid channels were analyzed separately for segmentation, morphological feature extraction, and deep learning applications.

#### Kidney tubule segmentation

Segmentation of kidney tubules from NoRI images was performed using a YOLOv8 instance segmentation model via the Ultralytics framework (Jocher et al., 2023). We trained the model as a single-class instance-segmentation model using both protein and lipid NoRI signals as input. Training images were divided into 640 × 640-pixel tiles. Model training was conducted with a batch size of 16 for up to 200 epochs, employing early stopping if no improvement was observed after 30 epochs. Data augmentation strategies included image flipping, rotation, adjustments to brightness and contrast, and affine transformations. Each channel was normalized by clipping intensities at the 1st and 99th percentiles and scaling to the maximum value. The resulting instance masks were used to delineate individual tubules and served as the basis for subsequent morphological analyses at the tubule level.

#### Tubule-type classification

Segmented tubules were classified as proximal, distal, or collecting based on antibody labeling. Anti-aquaporin-1 (AQP1), anti-uromodulin (UMOD), and anti-aquaporin-2 (AQP2) were used as specific markers for proximal, distal, and collecting tubules, respectively. Podocytes were identified using anti-podoplanin (T1α). Tubule identities inferred from marker staining were mapped to the corresponding NoRI structures and used for downstream analyses by tubule type.

#### Segmentation of subcellular and intratubular structures

Additional tissue structures—including nuclei, nucleoli, brush borders, tubular lumina, and lipid droplets—were segmented using individual U-Net convolutional neural networks (U-Net; Ronneberger et al., 2015). Ground-truth masks for each structure were manually annotated according to their morphology in NoRI protein and/or lipid images. The nuclei segmentation model used 640 × 640-pixel protein-channel images, while the nucleoli model was trained with 32 × 32-pixel paired protein and lipid images. Segmentation models for brush border, lumen, and lipid droplets were trained using 128 × 128-pixel paired protein and lipid channels. All U-Net models employed an encoder–decoder architecture with skip connections and were implemented in PyTorch (Paszke et al., 2019). Training used the Adam optimizer (Kingma & Ba, 2015; learning rate 1 × 10^−4^), binary cross-entropy loss, and a batch size of 8, with up to 200 epochs and early stopping after 30 epochs without improvement. Geometric and photometric augmentations were applied, and intensities were normalized by clipping at the 1st and 99th percentiles followed by scaling to the maximum. The checkpoint with the highest validation Dice score (Dice, 1945) was selected for subsequent analysis.

#### Quantification of NoRI morphological features

Morphological and biochemical features were calculated for each segmented tubule using the protein and lipid NoRI channels, in combination with masks for tubule, lumen, nuclei, nucleoli, brush border, and lipid droplets. Extracted features encompassed multiple aspects of tubule morphology. Brush-border organization was quantified by coverage, fraction, fragmentation, largest component fraction, and thickness. Radial organization of protein and lipid signals was assessed using apical-to-basal ratios, radial gradients, protein–lipid correlation, and radial roughness. Tubule and lumen geometry were described by quantitative metrics including lumen size and fraction, tubule radial roughness, concavity, solidity, epithelial thickness variability, eccentricity, and perimeter-to-area ratio. Protein and lipid abundance was measured across compartments such as tubule, lumen, nuclei, perinuclear regions, and brush border. Additional features included nuclear and nucleolar morphology, lipid-droplet abundance, tissue signal heterogeneity, and related structural parameters. Morphological features were determined individually for proximal, distal, and collecting tubules.

#### NoRI outcome endpoints

NoRI outcomes are animal means over sampled two-dimensional profiles of the contralateral kidney. The five endpoints shown in **Supplementary Figure 8D** use 31 animals. Models retain the animal as replicate. Displayed percentages divide the modeled effect and interval by the observed 9-month mean. Profile classification, density and area cannot be converted into total cells per kidney without additional sampling information.

### Deep-learning prediction of morphological age

#### Multimodal deep-learning model for morphological age prediction

A multimodal deep-learning model was developed to predict chronological age directly from NoRI morphology. Protein and lipid images were treated as two separate imaging modalities. Tissue images were divided into 320 × 320-pixel tiles using a sliding window with 50% overlap, corresponding to a stride of 160 pixels. Each tile was assigned the chronological age of the corresponding animal as the continuous regression target. Protein and lipid tiles were processed by separate ImageNet-pretrained (Russakovsky et al., 2015) ResNet-50 (He et al., 2016) encoders. The first convolutional layer of each encoder was modified to accept the corresponding single-channel NoRI modality. Encoder parameters were fine-tuned during training. Each encoder generated a 128-dimensional representation of its respective modality. The protein and lipid embeddings were subsequently combined using an attention-based multimodal fusion module (Vaswani et al., 2017). A learnable classification (CLS) token was concatenated with the protein and lipid modality tokens, and the resulting sequence was processed by a multi-head self-attention layer containing four attention heads. The attention module operated on 128-dimensional representations and included residual connections, layer normalization, and a feed-forward network. The resulting CLS representation was used as the fused multimodal representation. The fused representation was passed through a regression module consisting of a 128-dimensional fully connected layer, ReLU activation, dropout with a probability of 0.3, and a final regression layer producing a single continuous predicted-age value for each image tile. Attention weights were retained to assess the relative contribution of protein and lipid modality tokens to the fused representation. These weights were interpreted as measures of model attention to the two imaging modalities rather than as direct measures of biological importance.

#### Domain-adversarial batch correction

To reduce the influence of acquisition-related batch variation on the learned representations, domain-adversarial neural network (DANN) training (Ganin et al., 2016) was incorporated into the age-prediction model. Images belonged to two acquisition domains characterized by different imaging noise profiles. The fused multimodal representation was passed through a gradient-reversal layer and a domain-classification head that predicted the corresponding acquisition domain. The gradient-reversal operation encouraged the encoders and multimodal fusion module to learn representations informative for age prediction while reducing information predictive of imaging batch. Age prediction was optimized using mean squared error loss, whereas domain classification was optimized using cross-entropy loss. The combined objective was defined as L_total = L_age + 0.2 × L_domain, where L_age denotes the age-regression loss, and L_domain denotes the domain-classification loss. The strength of gradient reversal was progressively increased during training according to the domain-adversarial training schedule.

#### Model training and cross-validation

Models were trained using AdamW (Loshchilov & Hutter, 2019) with a learning rate of 3 × 10^−5^, weight decay of 1 × 10^−4^, and a batch size of 32 for up to 200 epochs. The model checkpoint with the lowest validation mean absolute error (MAE) was retained. Because checkpoints were selected on the validation animals, the reported errors are validation estimates rather than independent test results. Cross-validation was performed at the animal level to prevent image tiles from the same animal from appearing in both the training and validation sets. Of the imaged animals in each age group, three were reserved exclusively for training because these samples were also required for batch correction. The remaining animals (four or five per age group) participated in cross-validation. Ten cross-validation iterations were performed. In each iteration, one of the cross-validation animals from each age group was used for validation, while the remaining eligible animals together with the three training-only animals were used for model training. Across the ten iterations, each of the five cross-validation animals in the 18-, 21- and 25-month groups appeared in the validation set twice. Validation predictions were therefore made on animals not used to train that model. Predictions from each animal’s validation appearances were median-pooled to obtain the final validation prediction for that animal. The three training-only animals from each age group were excluded from downstream analyses requiring unbiased model predictions. Model performance was evaluated using mean absolute error (MAE) and the coefficient of determination (R²). Tile-level predictions were aggregated within animals using the median to obtain animal-level predicted morphological age.

#### Spatial mapping of predicted morphological age

The age-regression model generated an independent continuous age prediction for every 320 × 320-pixel NoRI tile. To visualize spatial heterogeneity in tissue aging, tile-level predictions were mapped back to their original coordinates within the tissue image. Because adjacent tiles overlapped by 50%, individual spatial locations could be represented in multiple tiles. For each spatial location, predicted ages from all overlapping tiles containing that location were averaged. This procedure generated a continuous spatial map of predicted morphological age across the imaged kidney tissue. These spatial predicted-age maps represent local model predictions derived from tissue morphology and should be distinguished from gradient-based activation maps such as Grad-CAM (Selvaraju et al., 2017). No Grad-CAM or equivalent pixel-level CNN activation method was used for the spatial age maps described here.

#### Tubule-specific morphological age prediction

To evaluate morphological aging at the level of individual renal tubules, the same multimodal age-regression framework was applied to segmented tubule-centered images. For each segmented tubule, a 320 × 320-pixel image was centered on the tubule. Pixels located outside the corresponding tubule mask were set to zero in both the protein and lipid channels, thereby restricting the model input to the morphology of the selected tubule. A single tubule age-prediction model was trained using proximal, distal, and collecting tubules together rather than training separate models for each tubule class. Predictions were subsequently stratified by tubule type for downstream analyses. Proximal and distal tubules were analyzed individually, whereas collecting tubules were retained during model training but were not evaluated as a separate age-acceleration group because of their smaller number.

#### Morphological age acceleration

Morphological age acceleration (Horvath, 2013) was defined as the difference between the morphological age predicted from NoRI and the age expected for an animal or tissue structure of the same chronological age. Expected predicted age was estimated using linear regression across the four chronological age groups: Expected morphological age = β_0_ + β_1_ × chronological age. Morphological age acceleration was then calculated as the residual:

#### Morphological age acceleration = predicted morphological age – expected morphological age

Positive residuals therefore indicate morphology predicted to be older than expected for chronological age, whereas negative residuals indicate morphology predicted to be younger than expected. Morphological age acceleration was evaluated for whole-tissue tile predictions and, separately, for predictions obtained from segmented tubule-centered images. Tubule-level analyses were performed for all tubules combined and separately for proximal and distal tubules.

#### Association of quantitative morphology with age and age acceleration

To identify interpretable morphological characteristics associated with kidney aging, quantitative NoRI features were compared with three age-related measures: chronological age, CNN-predicted morphological age, and morphological age acceleration. Although morphological features were initially calculated for individual tubules, statistical association analyses were performed at the animal level to avoid treating multiple tubules from the same animal as independent biological replicates. For each animal, individual-tubule feature values were aggregated using the median. Predicted morphological age and morphological age acceleration were likewise summarized at the animal level using median values. Associations between individual morphological features and chronological age, predicted morphological age, or morphological age acceleration were evaluated using Spearman rank correlation. Spearman’s correlation coefficient (p) and corresponding two-sided *P* values were reported. Differences in morphological features across the four chronological age groups (9, 18, 21, and 25 months) were evaluated using the Kruskal–Wallis test (Kruskal & Wallis, 1952). Statistical significance was defined as *P* < 0.05. Unless otherwise specified, reported *P* values for these exploratory morphology analyses were unadjusted for multiple comparisons.

#### Visualization of learned morphological representations

Uniform Manifold Approximation and Projection (UMAP) (McInnes et al., 2018) was used to visualize the high-dimensional representations learned by the multimodal age-prediction model. UMAP was used as a visualization method and not for statistical inference. The UMAP transformation was fitted exclusively using embeddings from the training dataset. Validation embeddings were subsequently projected into the fitted UMAP space without refitting the manifold, thereby preventing validation observations from influencing construction of the visualization space. The UMAP implementation was initialized with random_state = 10; other UMAP parameters were left at their default values. Validation embeddings were visualized in separate UMAP plots colored by chronological age and by CNN-predicted morphological age to illustrate the organization of the learned morphological representation with respect to both measured and model-predicted aging.

#### Integration of omics signatures with NoRI spatial features

Prediction of molecular abundance from NoRl images using convolutional neural networks and multiple-instance learning. To investigate spatial relationships between renal tissue morphology and molecular abundance, we developed a two-stage deep-learning framework integrating quantitative NoRl images with matched animal-level proteomic and metabolomic measurements. Candidate proteins and metabolites were prioritized based on their associations with chronological age and with quantitative NoRl-derived morphological and compositional features. For selected candidates, molecular-prediction models were trained within individual age groups to investigate relationships between tissue morphology and molecular abundance independently of broad chronological age differences. Quantitative NoRl protein and lipid images were divided into spatially indexed tiles of 320 × 320 pixels (132.48 × 132.48 µm), with 50% overlap, with each animal represented by a collection of image tiles matched to the corresponding molecular abundance measured in its contralateral kidney. Protein and lipid signals were assigned to the red and green input channels, respectively, with the third channel set to zero. lmage tiles were processed using an lmageNet-pretrained ResNet50 convolutional neural network fine-tuned for molecular abundance regression to generate morphological representations encoding protein and lipid image information. The CNN backbone generated 2,048-dimensional representations, which were standardized and reduced to 32 dimensions using principal component analysis (PCA) fitted on training data.

Tile-level embeddings were subsequently integrated using an additive multiple-instance learning (MlL) architecture, enabling prediction of animal-level protein or metabolite abundance from collections of tissue images. The MlL model incorporated learned sigmoid relevance gates and tile-specific contributions, allowing the predicted molecular abundance to be decomposed into contributions from individual image regions. Animal-level predictions were obtained by aggregating gated tile contributions with a learned bias term. Models were optimized using mean squared error (MSE) loss and the AdamW optimizer. The CNN was configured with a learning rate of 3 × 10^−4^, weight decay of 10^−4^, batch size of 32, and 50 training epochs. The MlL model used a learning rate of 10^−3^, weight decay of 10^-2^, dropout of 0.25, and up to 300 training epochs. During MlL training, up to 2,000 tiles were randomly sampled per animal, whereas all available tiles were used for validation, with performance evaluated using animal-level mean absolute error (MAE) and comparison with a training-mean baseline. Training and validation were separated at the animal level to prevent image tiles from the same specimen contributing to both datasets.

To investigate the spatial features associated with molecular predictions, learned tile contributions were mapped onto their original tissue coordinates, generating spatial representations of image regions contributing to predicted molecular abundance. Contributions from overlapping tiles were averaged to generate continuous spatial maps. Additional CNN activation maps were generated using Grad-CAM applied to ResNet50 convolutional layers 2–4, using gated MlL tile contributions as the attribution target, to visualize image features associated with individual molecular targets. Together, these approaches enabled the identification of spatially localized NoRl image patterns predictive of protein and metabolite abundance, despite molecular measurements being available only at the whole-animal level.

**Figures 6B,C** and **7B** compare, for each animal, the abundance predicted from NoRl images with the measured abundance, both relative to 9 months. Agreement is summarized by Spearman’s p across the plotted animals; where a *P* value is given, it uses the t approximation with n – 2 degrees of freedom.

## Statistical analysis

Analyses of NoRI features and image models used MATLAB, R, Python and GraphPad Prism v.11.1.0 (MA, US). Quantitative NoRI data, including the distributions of raw protein and lipid concentrations between groups, were compared using Kruskal–Wallis tests (Kruskal & Wallis, 1952), followed by Mann–Whitney *U* comparisons with Benjamini–Hochberg correction, as specified in the figure legends. Effect sizes were estimated as Hedges’ *g* or Cohen’s *d* (Cohen, 1988). Statistics specific to **Figures 6 and 7** were as follows. **Figure 6A**: orange outlines mark proteins and metabolites whose abundance correlated with age (Spearman’s p with age in months; Benjamini–Hochberg q < 0.05 across all 11,040 genes or all 269 metabolites). **Figures 6G and 7G**: for each molecule and NoRI feature (animal medians over all segmented objects), the standardized coefficient of the feature in an ordinary least-squares model of the molecule’s abundance with age group as a categorical covariate; dots mark nominal P < 0.05, and Benjamini–Hochberg *q* values were computed within the protein and metabolite families (**Figure 6G**) or within each block of molecules (**Figure 7G**). **Figures 7E and 7F**: age-adjusted Spearman correlations with KIM-1, computed as the Pearson correlation of ranks after removing each age group’s mean rank; program scores are the mean log_2_ abundance of their members, with KIM-1 excluded. Intervals in **Figure 7E** are the 2.5th–97.5th percentiles of 2,000 resamples of rats within age groups, and percentages in **Figure 7F** are the share of 1,000 such resamples in which a metabolite ranked among the ten strongest. Statistical significance was set at *P <* 0.05. Figures were drawn with matplotlib (Hunter, 2007) and GraphPad Prism.

## Acknowledgements

We thank Dr. Avik Mukherjee, Dr. Evangelos Gatzogiannis and Dr. Xili Liu for their help in maintaining the NoRI microscope.

## Author Contributions

*Research design and conceptualization:* WVT, MD, KP, MS, SO, RD, TS, DJG, MWK.

*Primary data collection:* WVT, MD, MS, KP, CM, HS, AG, RD, TS, BZ.

*Data analysis and interpretation:* WVT, MD, MS, KP, SO, BZ.

*Manuscript writing:* WVT, MD, MS, KP, SO, BZ, ETC, DJG, MWK.

## Data availability

Protein abundances for the 11,485 proteins in all 31 rats (log2, relative to the 9-month mean) are provided in **Supplementary Table 2**, and metabolite abundances for the 269 annotated features in the 31 metabolomes in **Supplementary Table 3**. The kidney RNA data analyzed here were published by Shavlakadze et al. (2026).

## Code availability

Code and materials used for image processing, together with the MATLAB and Python scripts used for statistical analysis, are available at: https://github.com/kirschnerlab/NoRI, https://github.com/HMS-IAC/nori, https://github.com/MDyakova/NORI_feature_extraction and https://github.com/MDyakova/NORI_image_segmentation.

