## Supplementary Figures 1-11 for "Quantitative imaging of native tissue architecture combined with deep multi-omics reveals the spatial biology of kidney aging"

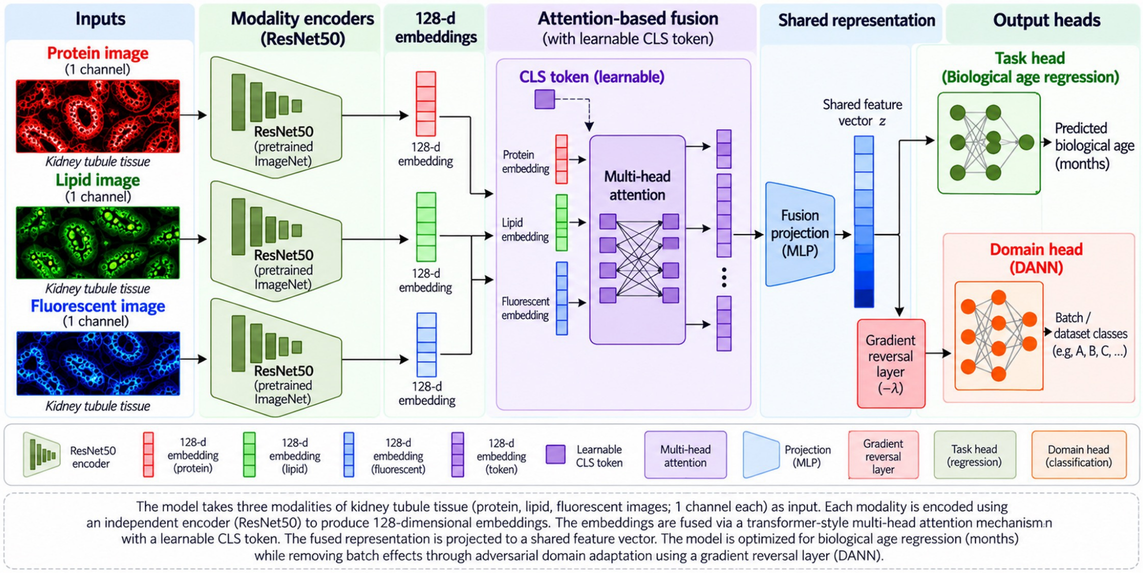


**Supplementary Figure 1. CNN model architecture.** The models in this study used the protein and lipid NoRI image layers; for future validation with targeted antibodies, an additional fluorescence layer can be added to the pipeline. The biological age regression in ‘Output heads’ gives the predicted ages shown in **Figure 2**.

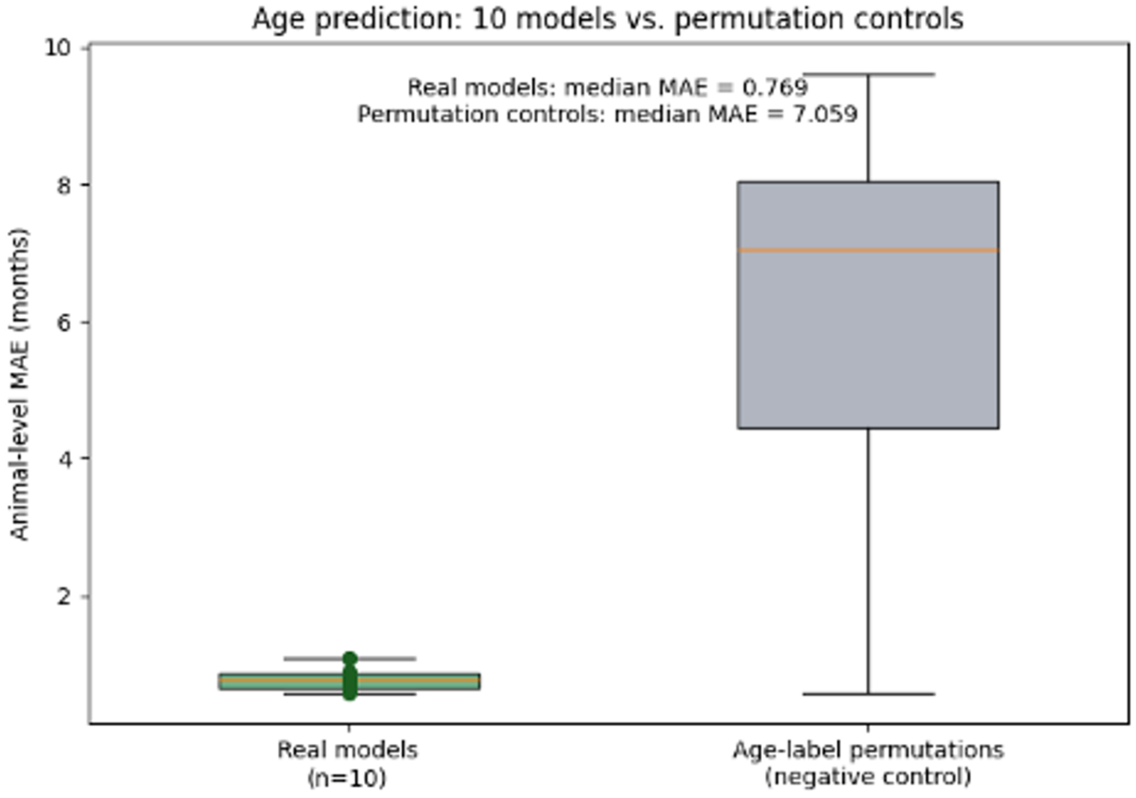


**Supplementary Figure 2. Validation of age prediction from NoRI images.**

Animal-level mean absolute error (MAE) of 10 separately trained CNN models (green), compared with the same models scored after shuffling age labels among the validation animals (gray; 10,000 shuffles per model). Median MAE was 0.77 months for the trained models and 7.06 months with shuffled labels. A few shuffles give low error by chance because each model was validated on only four animals.

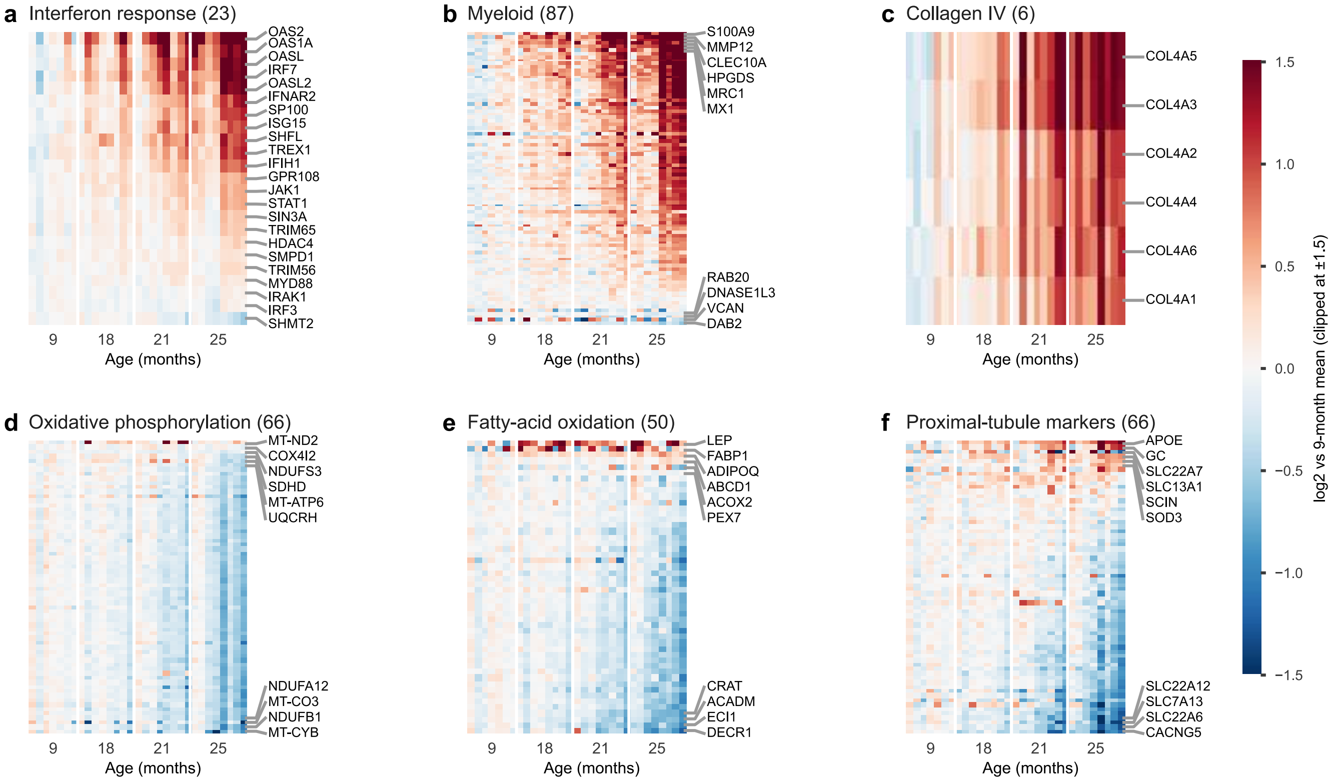


**Supplementary Figure 3. Protein-level structure of the programs.**

Member proteins (rows) of six programs in each rat (columns, ordered by age and then by KIM-1, as in Figure 4B), relative to each protein's 9-month mean. Rows are ordered by their change at 25 months. Small programs are fully labeled; in larger ones the six largest increases and four largest decreases are labeled. (A) Interferon response. (B) Myeloid. (C) Collagen IV. (D) Oxidative phosphorylation. (E) Fatty-acid oxidation. (F) Proximal-tubule markers.

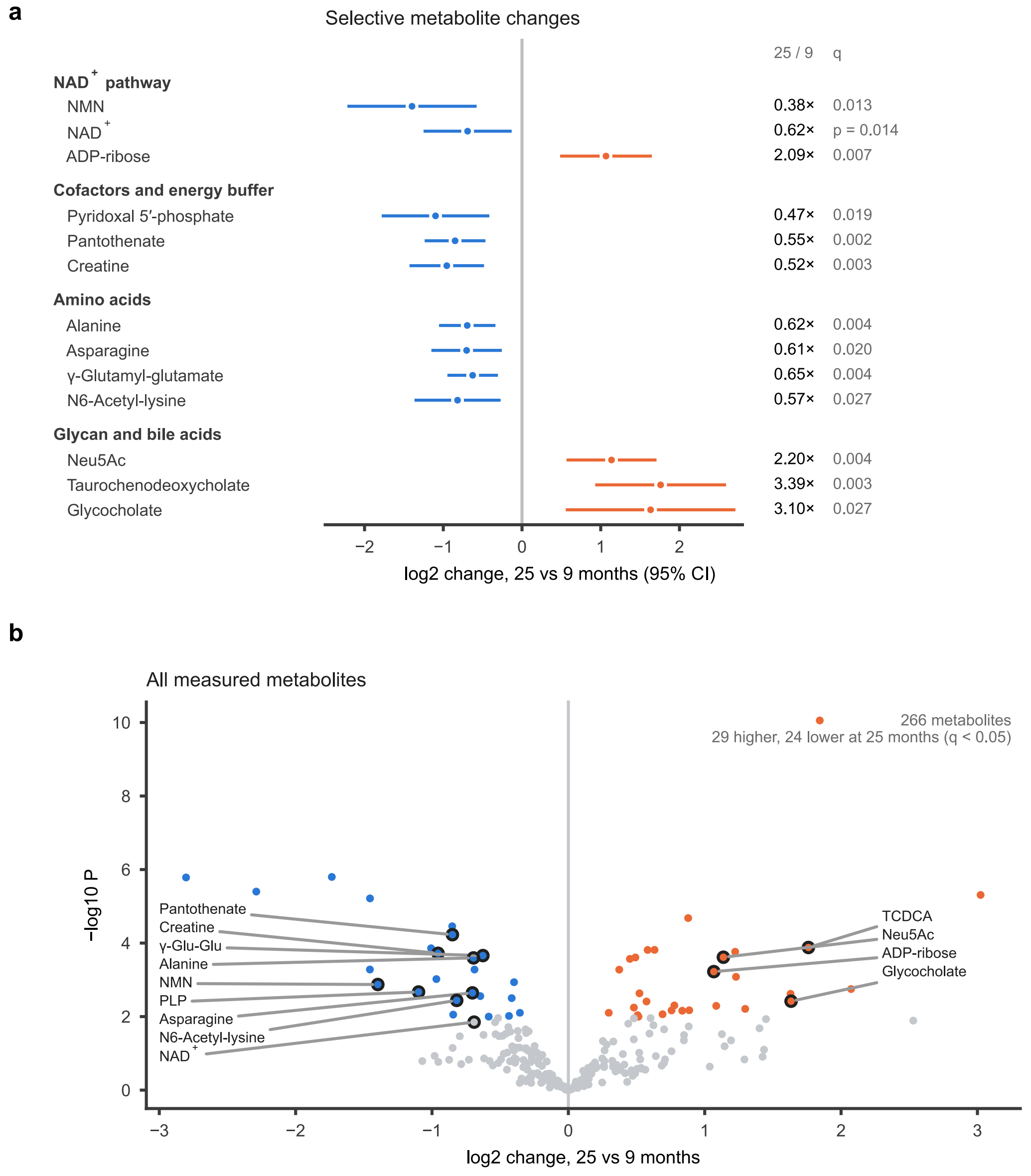


**Supplementary Figure 4. Selective metabolite changes.**

(A) Change between 25 and 9 months (log2, 95% CI) for selected metabolites, with fold change and Benjamini–Hochberg q across all 269 metabolites at right. (B) All 266 metabolites: change between 25 and 9 months against −log10 P; colored, Benjamini–Hochberg q < 0.05 across all 269 features (29 higher and 24 lower at 25 months; the summed NAD peaks have no q and are not colored); outlined, the metabolites in A. $NAD^+$ appears once, as the sum of its two peaks, and the two cyclic ADP-ribose features, which track $NAD^+$, are omitted. $NAD^+$ is the sum of its two chromatographic peaks and is shown with its nominal P. n = 8, 8, 7 and 8 rats at 9, 18, 21 and 25 months (one 21-month metabolome was excluded). Orange, higher at 25 months; blue, lower.

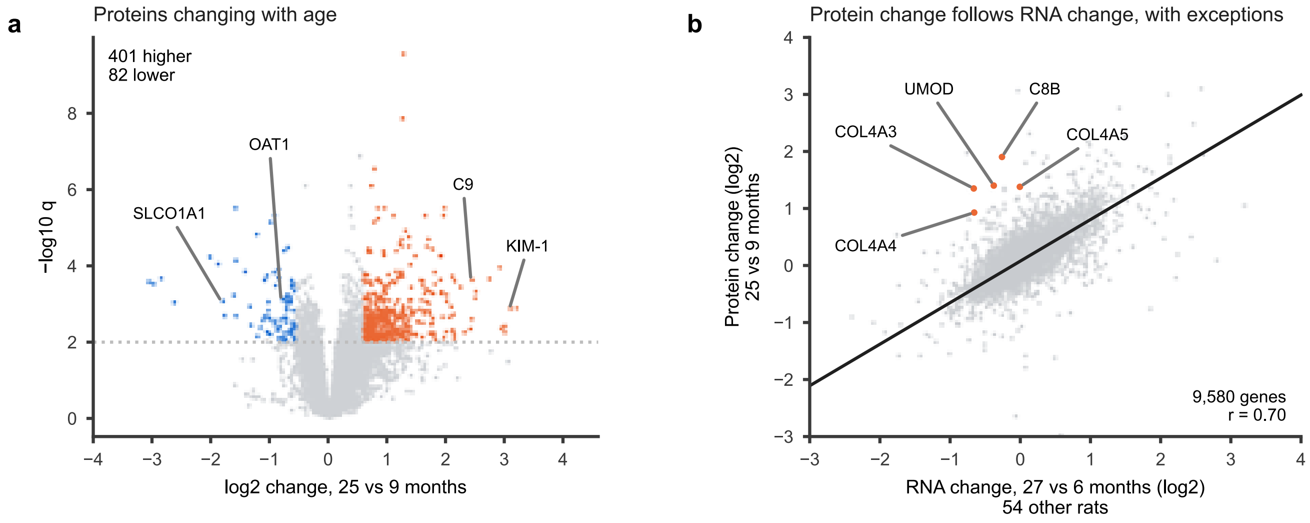


**Supplementary Figure 5. Proteins that change with age, and comparison with kidney RNA.**

(A) Change between 25 and 9 months for each gene against its significance (age-only model; Benjamini–Hochberg q). Colored genes have $q < 0.01$ and change more than 1.5-fold (401 higher, 82 lower); in all, 956 genes have $q < 0.01$. (B) Protein change (25 versus 9 months) against RNA change (27 versus 6 months) in kidneys of 54 other rats of the same colony (Shavlakadze et al., 2026), for 9,580 genes ($r = 0.70$; line, robust regression). The labeled extracellular proteins rise far more as protein than as RNA.

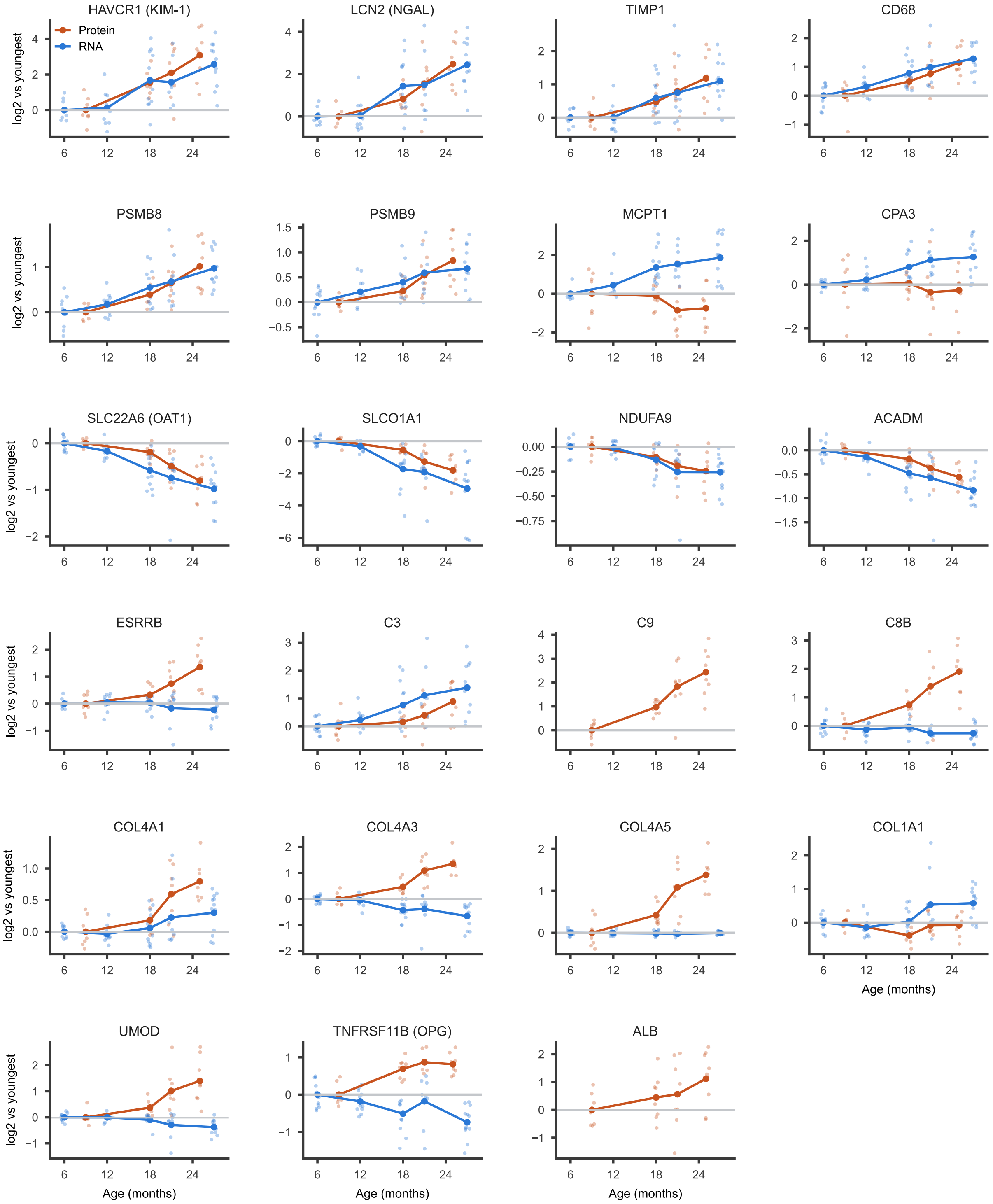


**Supplementary Figure 6. Protein and RNA trajectories of example genes.**

Protein (orange; this study, 9 to 25 months) and RNA (blue; 54 other rats, 6 to 27 months; Shavlakadze et al., 2026), each relative to its own youngest group. Small points are rats; lines are age-group means. C9 and albumin have no comparable kidney RNA data. Osteoprotegerin (TNFRSF11B) and ESRRB are each supported by a single unique peptide.

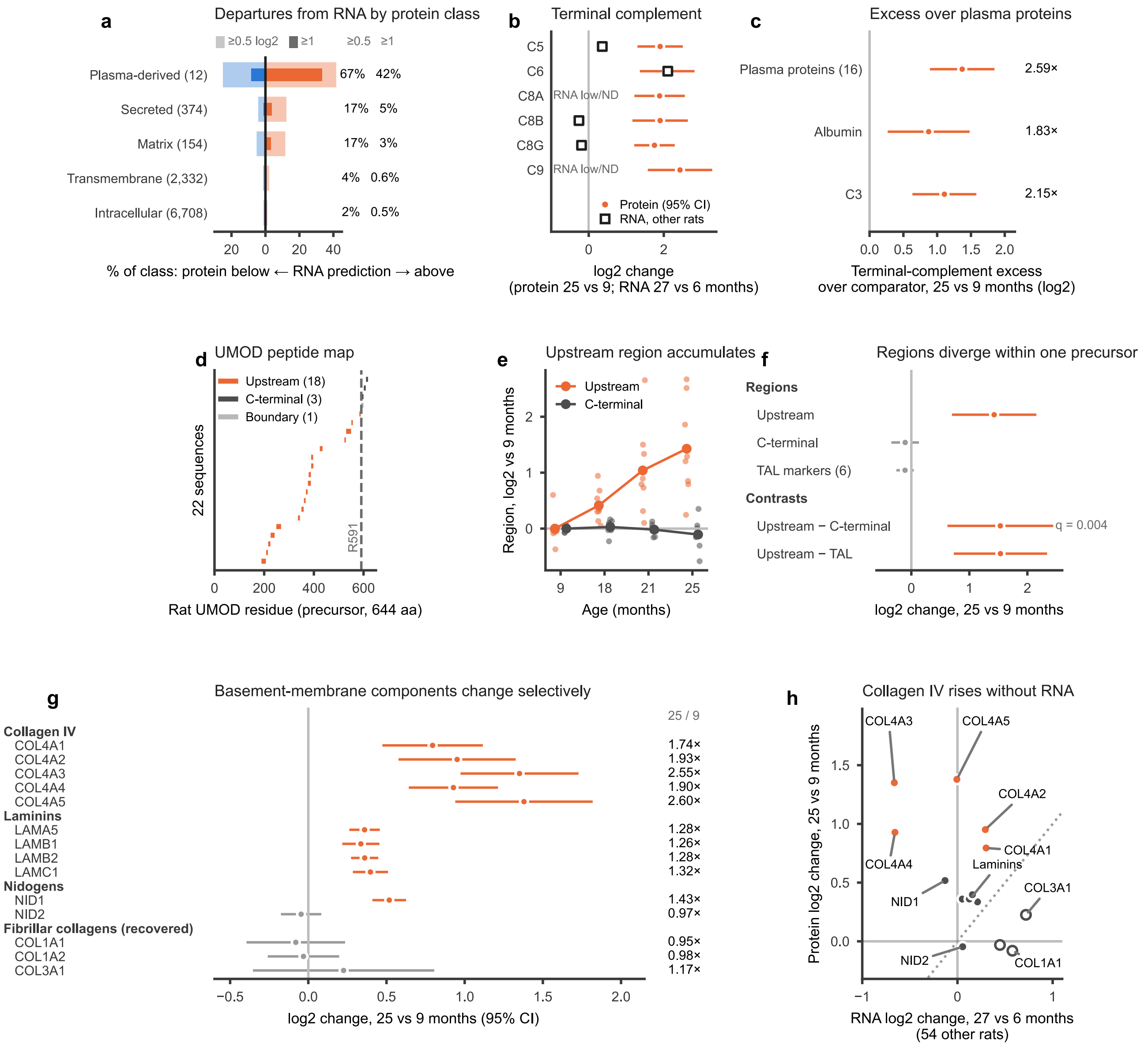


**Supplementary Figure 7. Extracellular proteins accumulate beyond their RNA.**

Protein changes are 25 versus 9 months in this study; RNA changes are 27 versus 6 months in kidneys of 54 other rats (Shavlakadze et al., 2026). (A) Share of each protein class whose protein change lies above (orange) or below (blue) the change predicted from its RNA by the fit in Supplementary Figure 5B, by at least 0.5 log2 (pale) or 1 log2 (dark); numbers at right are the percentages departing in either direction at each threshold. Plasma-derived proteins most often exceed their RNA, followed by secreted and matrix proteins. (B) Terminal complement proteins: protein change (points, 95% CI) and RNA change (squares); C8A and C9 RNA is low or absent in kidney. (C) Within each animal, terminal complement rises more than other plasma proteins, albumin or C3. (D) Positions of the 22 UMOD peptides on the rat UMOD precursor; peptides before and after R591 form the upstream and C-terminal regions. (E) The upstream region rises with age while the C-terminal region does not. (F) Regional contrasts and the thick-ascending-limb marker score (95% CI). (G) Basement-membrane and fibrillar matrix proteins (95% CI; numbers at right, fold change at 25 versus 9 months); collagen IV rises most, especially the α3 and α5 chains of the glomerular basement membrane, whereas recovered fibrillar collagens do not change. (H) Protein change against RNA change for the same proteins: collagen IV protein rises while its RNA does not. Orange, collagen IV; dark gray, laminins and nidogens; open circles, fibrillar collagens; dotted line, equal protein and RNA change.

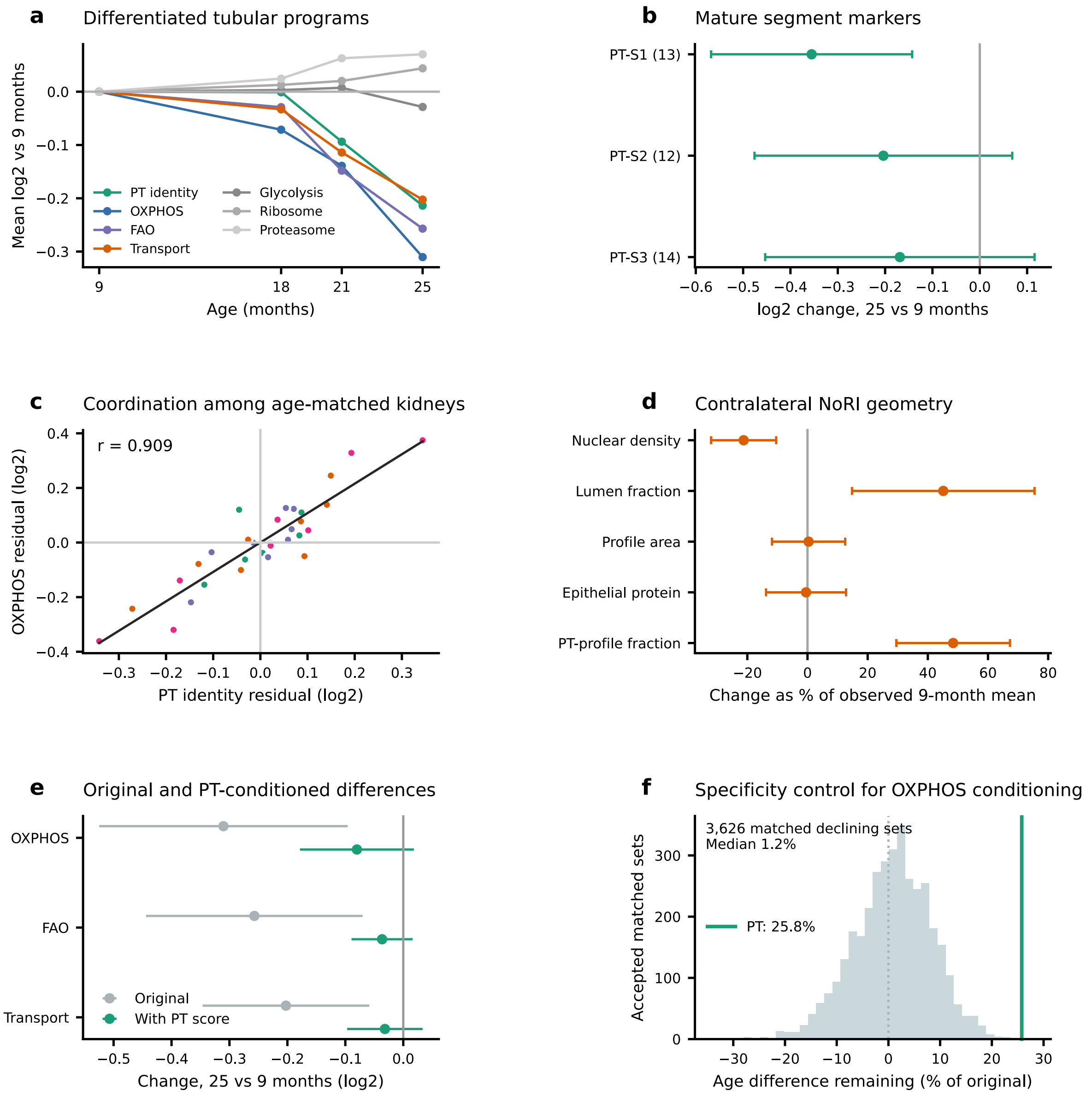


**Supplementary Figure 8. Proximal-tubule identity, metabolism and transport decline together.**

(A) Mean change of tubular programs and of general cellular machinery (glycolysis, ribosome, proteasome) with age. (B) Change in markers of the S1, S2 and S3 segments of the proximal tubule (25 versus 9 months, 95% CI). (C) Proximal-tubule identity and oxidative phosphorylation (OXPHOS) scores in each rat relative to its age group (r = 0.91). (D) Selected NoRI measurements from the other kidney (31 animals), as change from 9 months (percent of the 9-month mean, 95% CI). (E) OXPHOS, fatty-acid oxidation and transport changes before (gray) and after (green) accounting for proximal-tubule identity. (F) Control for E: random sets of other proteins that decline with age remove the OXPHOS age change at least as well as proximal-tubule identity does (green line), so the link shows coordination rather than a specific cause. n = 7, 8, 8 and 8 rats at 9, 18, 21 and 25 months.

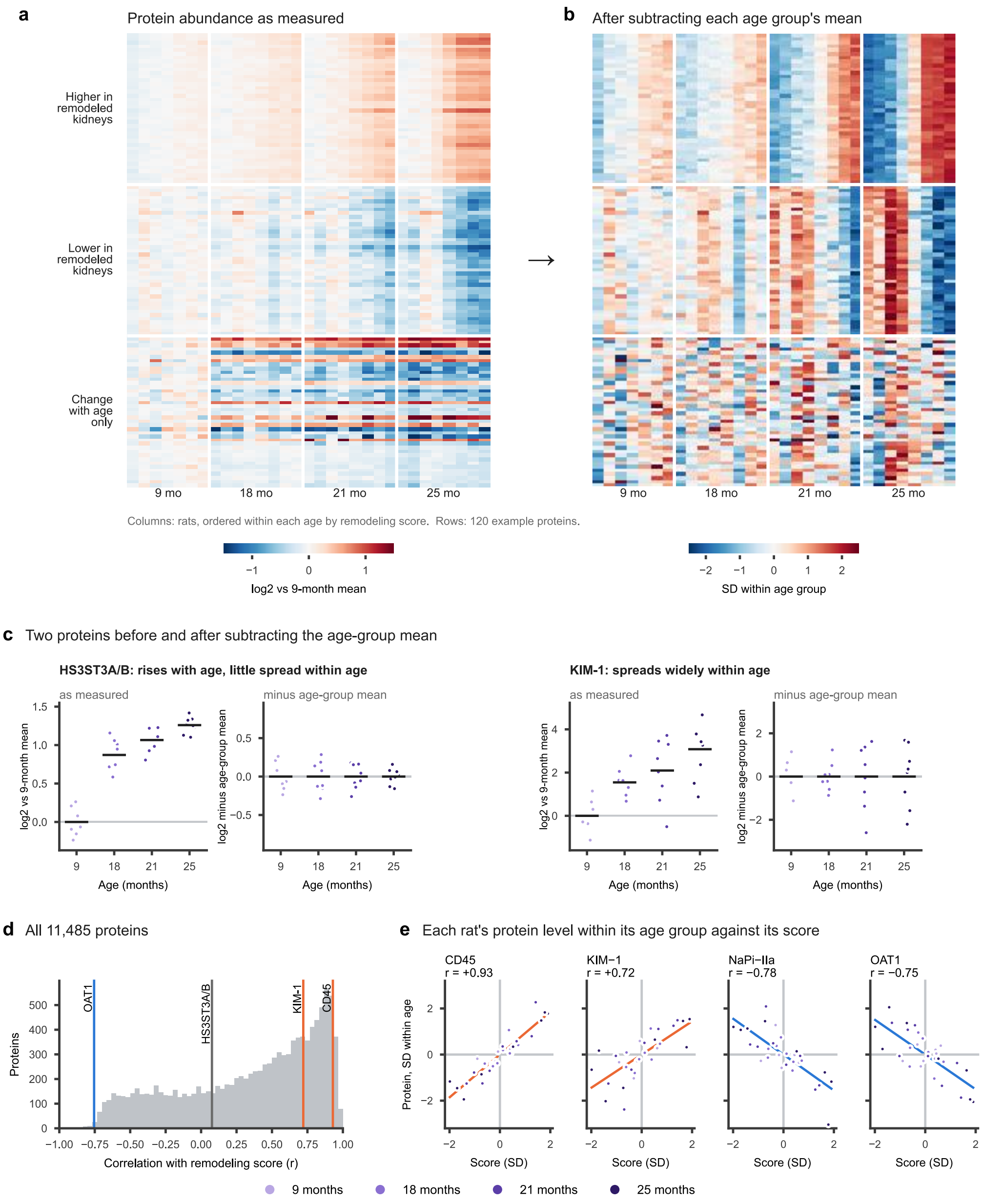


**Supplementary Figure 9. How the remodeling score is calculated.**

(A) Abundance of 120 example proteins as measured (log2 relative to the 9-month mean) in the 31 rats, grouped by age and ordered within each age by remodeling score: the 40 proteins most positively and the 40 most negatively correlated with the score, and 40 that change with age but barely track it. (B) The same proteins after subtracting each age group's mean and scaling each protein to unit standard deviation. The differences between ages disappear, and the coordinated differences among rats of the same age remain. The remodeling score is the first principal component of this matrix for all 11,485 proteins (36% of the variation within age groups), oriented so that higher CD45 is positive. (C) HS3ST3A/B and KIM-1 before and after subtracting the age-group mean, on the same scale: HS3ST3A/B changes with age but hardly varies within age, whereas KIM-1 varies widely among rats of the same age. (D) Correlation of each protein's within-age values with the score. (E) Within-age values of four proteins against the score, two at each end; lines are least-squares fits. n = 7, 8, 8 and 8 rats at 9, 18, 21 and 25 months.

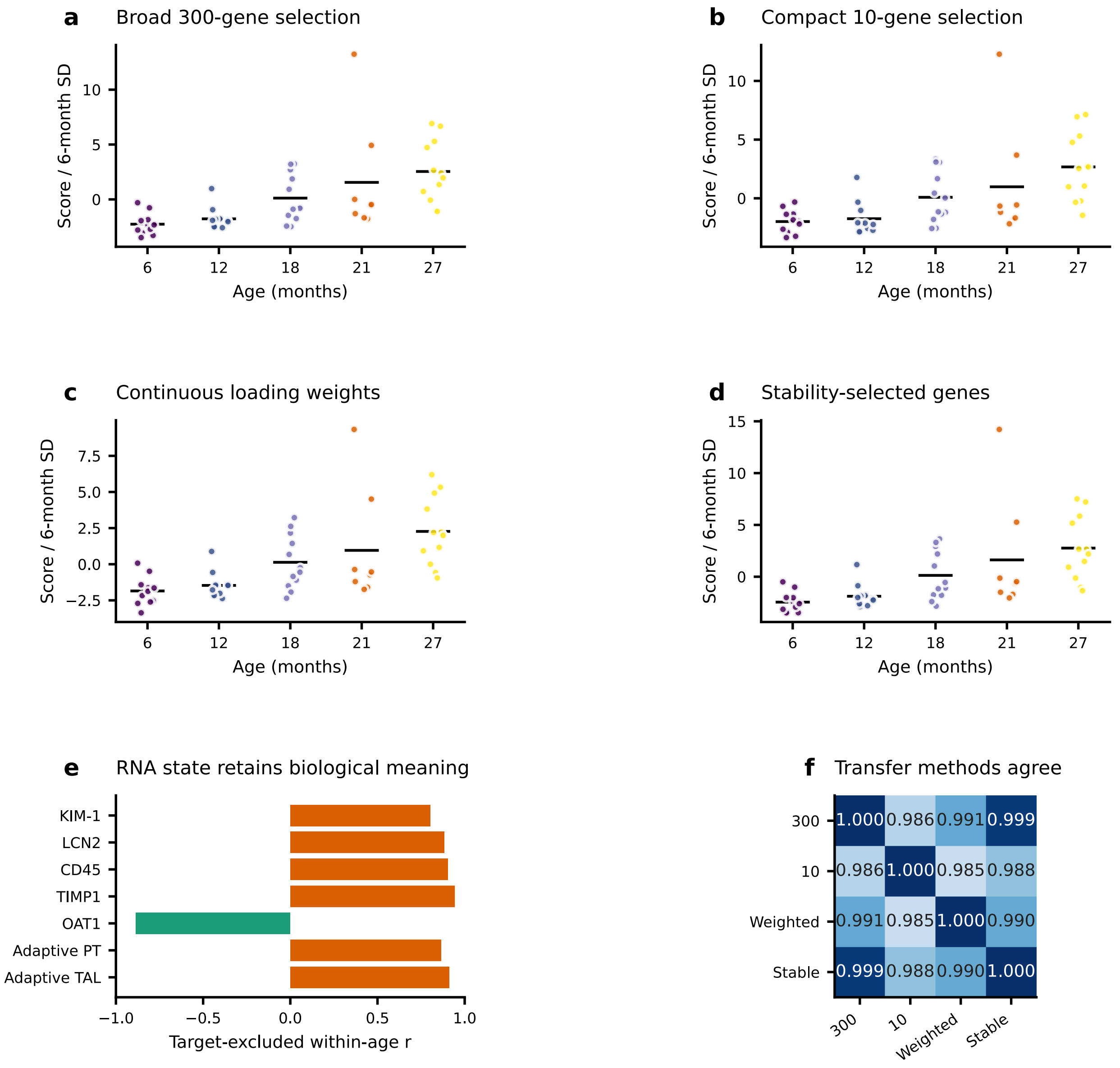


**Supplementary Figure 10. The RNA remodeling score does not depend on how it is built.**

(A–D) RNA remodeling score in 54 rats built four ways: the 300 top and 300 bottom proteins with equal weights (A; used in Figure 5E, where it is shown relative to 6-month rats), a compact version with 10 top and 10 bottom genes (B), continuous weights (C) and proteins selected in at least 60% of resamples (D). Points are rats; bars are age-group means. (E) Correlation of the continuous-weight score with marker transcripts among rats of the same age, each marker left out of the score. (F) The four scores agree closely ($r \geq 0.985$).

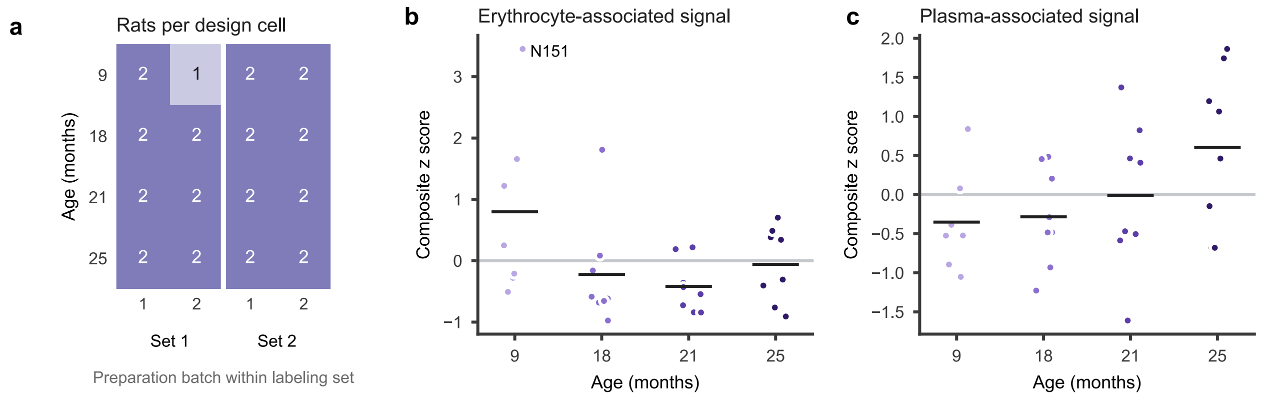


**Supplementary Figure 11. Study design balance and blood-derived signal.**

(A) Number of rats in each combination of age, TMT labeling set and preparation batch. One 9-month rat with a visibly bloody preparation was excluded. (B, C) Erythrocyte (B) and plasma (C) protein scores (Methods). Points are rats; bars are age-group means. N151 has the highest erythrocyte score and was kept in all analyses.
