## Supplementary Table 1 for "Quantitative imaging of native tissue architecture combined with deep multi-omics reveals the spatial biology of kidney aging"

**Supplementary Table 1.** Features derived from NORIGAMI segmentation of the rat kidney. The 51 protein, lipid and morphological features are further represented across the three tubule types measured in the present study: proximal tubules, distal tubules, and collecting ducts, resulting in 153 quantifiable features in our dataset.

| Original label | Biological Equivalent Label | Category | What it measures | High values represent |
|---|---|---|---|---|
| tb_size | Tubule cross-sectional area | Size | Area of the tubule in the section | Larger tubule cross-section |
| protein_level | Protein concentration (whole tubule) | Protein | Mean protein concentration across the tubule | More protein-rich material |
| lipid_level | Lipid concentration (whole tubule) | Lipid | Mean lipid concentration across the tubule | More lipid-rich material |
| protein_level_nuclei | Nuclear protein | Protein | Mean protein concentration inside nuclei | Protein-rich nuclei |
| protein_level_around_nuclei | Perinuclear protein | Protein | Mean protein concentration in a ring around each nucleus | More perinuclear protein |
| lipid_level_around_nuclei | Perinuclear lipid | Lipid | Mean lipid concentration in a ring around each nucleus | More perinuclear lipid |
| protein_level_lumen | Luminal protein | Protein | Mean protein concentration inside the lumen | Protein-rich lumen (e.g. casts) |
| lipid_level_lumen | Luminal lipid | Lipid | Mean lipid concentration inside the lumen | Lipid-rich lumen |
| lumen_size | Lumen area | Size | Area of the lumen | Dilated lumen |
| protein_level_bb | Brush-border protein | Protein | Mean protein concentration in the brush border | More brush-border protein |
| lipid_level_bb | Brush-border lipid | Lipid | Mean lipid concentration in the brush border | More brush-border lipid |
| bb_size | Brush-border area | Brush border | Absolute area of brush border | More brush border |
| protein_level_tb_conn | Inter-tubular protein | Protein | Mean protein concentration in the tubule-connection mask | More inter-tubular protein |
| lipid_level_tb_conn | Inter-tubular lipid | Lipid | Mean lipid concentration in the tubule-connection mask | More inter-tubular lipid |
| tb_conn_size | Inter-tubular region area | Size | Area of the tubule-connection mask | Larger inter-tubular region |
| black_holes | Signal voids / vacuoles | Morphology | Amount of very low-signal spaces inside the tubule | More/larger dark voids |
| nuclei_number | Nuclei per tubule | Nuclear | Count of nuclei | More nuclei |
| nuclei_area_mean | Nuclear size | Nuclear | Mean area of nuclei | Larger nuclei |
| nucleolus_number_in_nuclei | Nucleoli per nucleus | Nuclear | Number of nucleoli within nuclei | More nucleoli |
| nucleolus_area_in_nuclei | Nucleolar area | Nuclear | Area occupied by nucleoli within nuclei | Larger nucleoli |
| protein_delta | Protein compartment contrast | Protein | Difference in protein concentration between two | Larger protein difference |
| lipid_delta | Lipid compartment contrast | Lipid | Difference in lipid concentration between two compartments | Larger lipid difference |
| lipid_droplet_size | Lipid droplet size | Lipid droplets | Typical size of lipid droplets | Larger droplets |
| body_size | Cell body area | Size | Area of the legacy 'body' mask | Larger body region |
| lipid_droplet_level | Lipid droplet content | Lipid droplets | Amount and/or intensity of lipid-droplet signal | More or denser droplets |
| bb_size_k | Brush-border fraction | Brush border | Brush-border area normalised to a reference area | More brush border relative to tubule |
| lumen_size_k | Lumen fraction | Size | Lumen area as a fraction of tubule area | Relatively larger lumen |
| area_n | Total nuclear area | Nuclear | Nuclear area in the legacy pipeline | More nuclear area |
| nucleolus | Nucleoli | Nuclear | Legacy nucleolus count or area | More/larger nucleoli |
| area_nc | Nucleolar area | Nuclear | Legacy nucleolar area | More nucleolar area |
| robust_cv_protein | Protein heterogeneity | Protein | Patchiness of protein within the tubule | Uneven, patchy protein |
| robust_cv_lipid | Lipid heterogeneity | Lipid | Patchiness of lipid within the tubule | Uneven, patchy lipid |
| brush_border_coverage | Brush-border coverage | Brush border | Fraction of the luminal surface covered by brush border | Continuous brush border along the lumen |
| brush_fraction | Brush-border fraction of epithelium | Brush border | Brush-border area as a fraction of epithelial area | More brush border relative to epithelium |
| brush_fragmentation | Brush-border fragmentation | Brush border | Number of separate brush-border fragments per length of | Broken, fragmented brush border |
| largest_brush_component_fraction | Brush-border continuity | Brush border | Share of brush border in its largest continuous piece | One continuous brush border |
| brush_thickness | Brush-border thickness | Brush border | Median thickness of the brush border | Taller brush border |
| protein_apical_basal_ratio | Apical:basal protein ratio | Polarity | Protein near the lumen relative to protein near the basement | Protein enriched apically |
| lipid_apical_basal_ratio | Apical:basal lipid ratio | Polarity | Lipid near the lumen relative to lipid near the basement | Lipid enriched apically |
| protein_radial_gradient | Apical-to-basal protein gradient | Polarity | How consistently protein rises or falls from lumen to basement | Protein increases towards the basal side |
| lipid_radial_gradient | Apical-to-basal lipid gradient | Polarity | How consistently lipid rises or falls from lumen to basement | Lipid increases towards the basal side |
| protein_lipid_radial_correlation | Protein–lipid co-polarization | Polarity | Whether protein and lipid peak in the same zones across the | Protein and lipid co-organised |
| radial_roughness_cv | Lumen irregularity | Morphology | Unevenness of the lumen outline | Irregular lumen |
| concavity_fraction | Lumen indentation | Morphology | How much the lumen outline folds inwards | Deeply indented lumen |
| tubule_radial_roughness_cv | Tubule outline irregularity | Morphology | Unevenness of the outer tubule outline | Wavy, irregular tubule |
| tubule_concavity_fraction | Tubule indentation | Morphology | How much the outer tubule outline folds inwards | Deeply indented tubule |
| tubule_solidity | Tubule compactness | Morphology | How completely the tubule fills its convex outline | Compact, convex tubule |
| epithelial_thickness_cv | Epithelial thickness variability | Morphology | Variation in epithelial height around the tubule | Uneven thinning/thickening |
| tubule_eccentricity | Tubule elongation | Morphology | How elongated the tubule cross-section is | Elongated tubule |
| tubule_perimeter_area_ratio | Tubule outline complexity | Morphology | Outline length relative to tubule area | More convoluted or smaller tubule |
